# UTRGAN: Learning to Generate 5’ UTR Sequences for Optimized Translation Efficiency and Gene Expression

**DOI:** 10.1101/2023.01.30.526198

**Authors:** Sina Barazandeh, Furkan Ozden, Ahmet Hincer, Urartu Ozgur Safak Seker, A. Ercument Cicek

## Abstract

The 5’ untranslated region (5’ UTR) of mRNA is crucial for the molecule’s translatability and stability, making it essential for designing synthetic biological circuits for high and stable protein expression. Several UTR sequences are patented and widely used in laboratories. This paper presents UTRGAN, a Generative Adversarial Network (GAN)-based model for generating 5’ UTR sequences, coupled with an optimization procedure to ensure high expression for target gene sequences or high ribosome load and translation efficiency. The model generates sequences mimicking various properties of natural UTR sequences and optimizes them to achieve (i) up to 5-fold higher average expression on target genes, (ii) up to 2-fold higher mean ribosome load, and (iii) a 34-fold higher average translation efficiency compared to initial UTR sequences. UTRGAN-generated sequences also exhibit higher similarity to known regulatory motifs in regions such as internal ribosome entry sites, upstream open reading frames, G-quadruplexes, and Kozak and initiation start codon regions. In-vitro experiments show that the UTR sequences designed by UTRGAN result in a higher translation rate for the human TNF-*α* protein compared to the human Beta Globin 5’ UTR, a UTR with high production capacity.

## 1 Introduction

RNA-based therapeutics necessitate both tunability and a long-lasting profile after administration to the body. To achieve this, optimization of both mRNA molecules and carriers are crucial to enhance their stability and promote tissue-specific tropism [63]. The level of RNA and protein expression resulting from mRNA therapeutics plays a critical role in various applications such as protein replacement therapies, genome engineering, genetic reprogramming studies, as well as vaccination and cancer immunotherapies [55]. In specific protein replacement therapies for conditions like hemophilia A and cystic fibrosis exemplify the significance of targeted mRNA expression levels. For instance, Chen et al. demonstrated that to attain the required factor VIII expression in mice using mRNA-laden lipid nanoparticles (LNPs), a concentration of 2 mg mRNA per kilogram of mouse body weight is necessary [15]. Similarly, a study focusing on cystic fibrosis treatment found that a dosage of 0.1 mg/kg/day for two consecutive days was sufficient to restore the function of the CFTR gene in CFTR-knockout mice [54]. A phase 1 study addressing transthyretin amyloidosis utilized a dosage of 0.3 mg/kg to patients, resulting in the expression of Cas9 and the delivery of its single guide RNA to knock out the transthyretin gene, representing a potential treatment for this disease [26]. Conversely, vaccination studies for viruses like Zika [51], Covid [24, 65], and influenza [20], or tumor-associated antigen expression in cancer immunotherapies [56], require a lower dosage of approximately 0.002-0.02 mg/kg injection to induce immunity against these viruses and tumor cells. Thus, designing stable RNA molecules and being able to control the expression levels are desirable [2, 11, 58].

Optimization of the 5’ UTR is a preferable approach to control the stability and expression level of the mRNA, because it has been shown to be an important region in the sequence with regards to these features [39] [60]. Although UTR is not translated into the protein, it plays a crucial role in regulating the translation process because it contains the ribosome binding site (RBS) through which the ribosome attaches to initiate the translation. For this reason, many UTR sequences have been patented and used in the design of gene circuits [43, 67].

The standard approach for optimizing UTR sequences is to introduce variants in existing target sequences and evaluate their effectiveness using simple algorithmic approaches. Von Niessen et al. construct a library of 3’ UTR sequences from the human genome and identify 3’ UTR sequences that improve transcript stability [67] [47] [66]. Then, they generate synthetic sequences using a genetic algorithm based on sequence characteristics such as GC content, k-mer frequency, and free energy. The generated sequences are then selected with respect to their predicted translation efficiency [13]. Similarly, Lu et al. identify 5’ UTRs that affect protein expression and discuss methods for pairing them [43]. Studies also identify 3’ and 5’ UTRs for higher gene expression, and the samples are selected from naturally abundant mRNA sequences in Human tissues [10]. More recently, Chu et al. used a predictive language model paired with random mutation for designing a library of 5’ UTRs with high translation efficiency [17]. These models use machine learning algorithms only to predict sequence features and use those features to evaluate the generated samples. The potential of generative modeling is not fully utilized for designing and optimizing UTR sequences.

Several studies employ Generative Adversarial Networks (GANs) to generate sequences with similar characteristics to the natural DNA or RNA sequences [40, 69]. RNAGEN is a framework that enables the generation and optimization of synthetic piRNA sequences with some desired properties, such as binding to target proteins [50]. Zirmec et al. propose ExpressionGAN, a framework based on GANs for generating 1 kb-long regulatory DNA sequences (promoter, 5’UTR, 3’UTR, and terminator) [78, 79]. Using a model that predicts yeast gene expression, they optimize the full regulatory sequence considered (not just UTR) [77]. The model used here allocates a relatively short and fixed length for generated regulatory regions (250 bp for 5’ UTR and 1 kbp for the entire sequence), which would not fit many human genes with larger regulatory sequences, such as the MECP2 gene, where the length of the 3’ UTR extends over 8 kbs [18]. In another study, Castillo-Hair et al. discuss the potential of using machine learning methods along with predictive models for optimizing 5’ UTR sequences [14]. Linder et al. use predictive models for filtering generated *E. coli* promoter sequences and Wang et al. optimize DNA sequences generated using activation maximization for functional proteins [40, 69]. To the best of our knowledge, there is no prior work based on generative models that are tailored for generating and optimizing 5’ UTRs with respect to various metrics.

In this work, we propose UTRGAN, the first GAN-based pipeline for generating novel human 5’ UTR sequences and optimizing them to yield higher Mean Ribosome Load (MRL), Translation Efficiency (TE), and higher gene expression. Our model generates 5’ UTR sequences that can be attached to any natural or synthetic gene of interest and can generate UTR sequences with variable lengths. UTRGAN optimizes the gene expression of any desired human gene solely based on the coordinates of the TSS and the 5’ UTR regions. This enables gene expression optimization (or other desired targets as described below). UTRGAN can generate 5’ UTR sequences without modifying the target DNA/RNA sequence it is attached to. Our generated sequences resemble the natural 5’ UTRs with respect to the distribution of various important characteristics, such as GC content, k-mer distance, and Minimum Free Energy (MFE) [62]. Furthermore, the optimization procedure enables the generation of optimized sequences for higher MRL, TE, and mRNA abundance. By using multiple sequences for optimization, we show that we can increase the predicted MRL, mRNA expression by 53% and by 61%, respectively, compared to the initial values for the designed sequences. In addition, optimization for TE results in up to a 34-fold increase in the average predicted value. Both results show that the optimization procedure works as intended. Depending on the application, our model is able to optimize the generated 5’ UTRs for a specific target DNA sequence or for a set of DNA sequences. The optimization for a single gene of interest increases the expression 2.2-fold on average and can result in up to 32 times higher expression for the best synthetic 5’ UTR. We further analyze our sequences in terms of their similarity to known regulatory motives, including Internal Ribosome Entry Site (IRES) and Upstream Open Reading Frame (uORF) sequences, and demonstrate that sequences generated and optimized using UTRGAN maintain important motives and regulatory elements found in natural sequences. We also show that these motives are much less conserved in the sequences generated by other approaches, even if they have high predicted MRL values, suggesting that they might not be functional. We also conduct in-vitro experiments and demonstrate that UTRGAN-generated 5’ UTR sequences indeed yield a higher translation rate compared to the natural human *β*-globin 5’ UTR when attached to the TNF-*α* gene.

UTRGAN’s ability to generate and optimize 5’ UTRs for any given gene sequence will be the key enabler for the genetic circuit design. We think that UTRGAN will pave the way for mRNA-based therapeutics in the biotech industry for any application that requires gene expression control, such as cancer immunotherapy.

## 2 Results

### 2.1 Overview of UTRGAN

The UTRGAN model is a deep generative adversarial network (GAN) that learns to generate 5’ UTR sequences with characteristics similar to those of natural ones. This model is a variant of the WGAN-GP architecture [29] for one-hot encoded DNA sequence. The model provides improved training over the original Convolutional GAN (DCGAN) models [29,53] and is less prone to overfitting than the original WGAN architecture [4].

Figure 1A shows the overview of the architecture. The input is a noise vector. The generator and the critic are convolutional neural networks trained together. The generator upsamples the input vector using transpose-convolutions to generate 5’ UTR sequences, whereas the critic uses convolutions and dense layers to distinguish natural and synthetic 5’ UTRs. Based on this feedback, the generator learns to generate more natural-like 5’ UTRs. The optimization pipeline is shown in Figure 1B. We use the guidance of off-the-shelf deep-learning models for optimization. To optimize the gene expression of a target RNA sequence, we use Xpresso [1], which predicts the expression of a given sequence, including the UTR region. To optimize the MRL of a UTR sequence, we use FramePool [33], which predicts this value based on the UTR sequence only, independently from the RNA sequence it is attached to. Similarly, we use MTrans [76] to predict the translation efficiency of a UTR sequence. We perform optimization by updating the initial input to the GAN model, which generates the UTR sequence until the generated sequence converges to its maximum predicted value independently from other sequences in its batch. That is, we apply gradient ascent on the generator’s input with respect to the feedback from the target feature predictor (either MRL, translation efficiency, or gene expression).

**Fig. 1:**
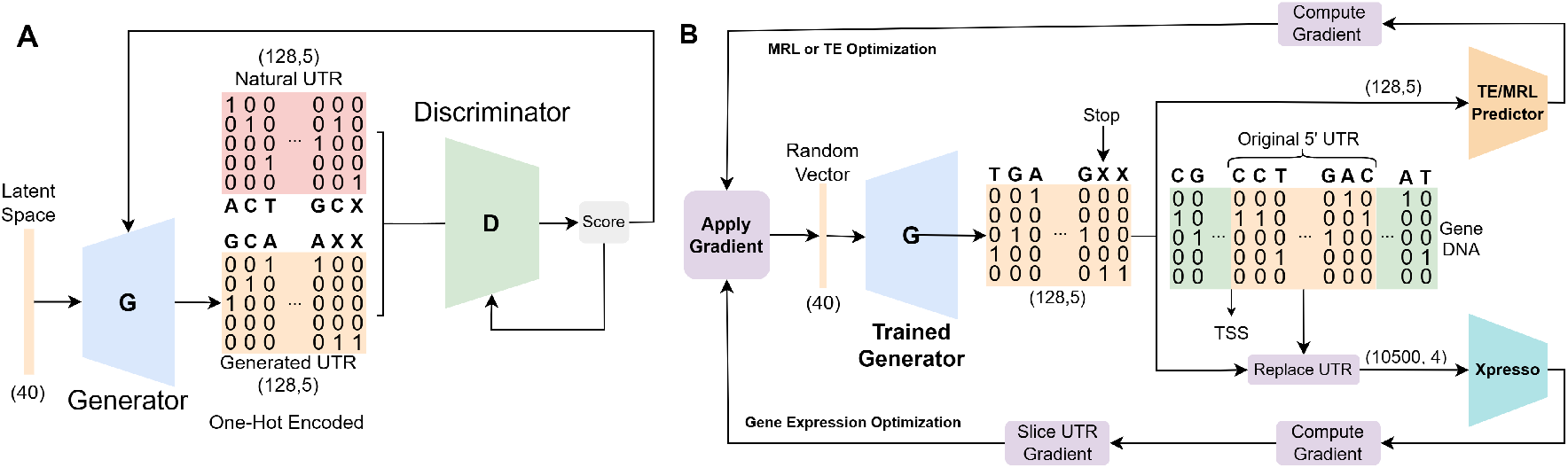
Generative and optimization architectures. Shown on panel **A** is the training phase of the GAN model. The feedback resulting from the competition between the critic and the generator updates the weights of both components. The panel **B** shows the optimization procedure. The GAN is used to generate samples, and the input noise is updated using the Gradient Ascent algorithm to increase the predicted MRL, TE, or mRNA abundance of the generated 5’ UTR sequences. The gene expression optimization procedure (Xpresso model) requires attaching the generated 5’ UTR to a DNA sequence to get feedback. We slice and apply the relevant gradient to update the input to generate the UTR sequence (the random vector). The MRL and TE optimization require only the UTR sequence as their input.

In the following subsections, we perform various analyses to show that generated 5’ UTR sequences resemble natural 5’ UTRs and compare our performance with other approaches. The only generation tool is our approach, UTRGAN. Thus, the generated sequences are always given by UTRGAN. These generated sequences are optimized by our approach, and we call such sequences UTRGAN-optimized sequences (expression, MRL, or TE as target feature). We also compare our approach with a randomized algorithm to optimize sequences, Optimus 5-Prime. We truncate our generated sequences to the desired input length for this method, which is 50 bps, and use them as the initial input sequences for this method. We call the sequences optimized by this method Optimus 5-Prime-optimized from now on.

### 2.2 Levenshtein distances are similar in natural and generated UTRs

A metric to measure the similarity across two sets of sequences is to use the distribution of the distances to the closest sequences in a target set (i.e., natural UTR set). Levenshtein distance is the minimum number of single-character edits (insertions, deletions, or substitutions) required to change one word into the other [38]. For this test, we compare the following sets: (i) UTRGAN-generated sequences (no optimization, n = 1204), (ii) 1024 UTRGAN-generated and MRL-optimized sequences (n = 1024), and (iii) sequences generated and MRL-optimized by Optimus 5-Prime [33] (n =1024).

We use the natural 5’ UTR dataset (n = 33,250) as the target set. In other words, we find the distribution of distances from the sequences in the above-mentioned sequence sets to the closest sequence in the natural UTR dataset with respect to the Levenshtein distance. This gives us three distributions, one for each of the following sequence classes: UTRGAN-generated, UTRGAN-optimized, and Optimus-5-Prime-optimized sequences. The same procedure is performed for the natural UTR set against itself to obtain the baseline distribution. There are a few natural sequences with anomalously lower distances to the natural UTR set, and we discard those in the plots.

Figure 2A shows that the UTRGAN-generated sequences and the natural sequences have almost identical distributions. We also observe that the UTRGAN-optimized sequences maintain their closeness to the natural sequences while also maintaining the wide sequence length range (See Supplementary Figure 5). On the other hand, sequences optimized using Optimus 5-Prime have a fixed 50bps length by design and they are less diverse with respect to their distances to natural UTR sequences.

**Fig. 2:**
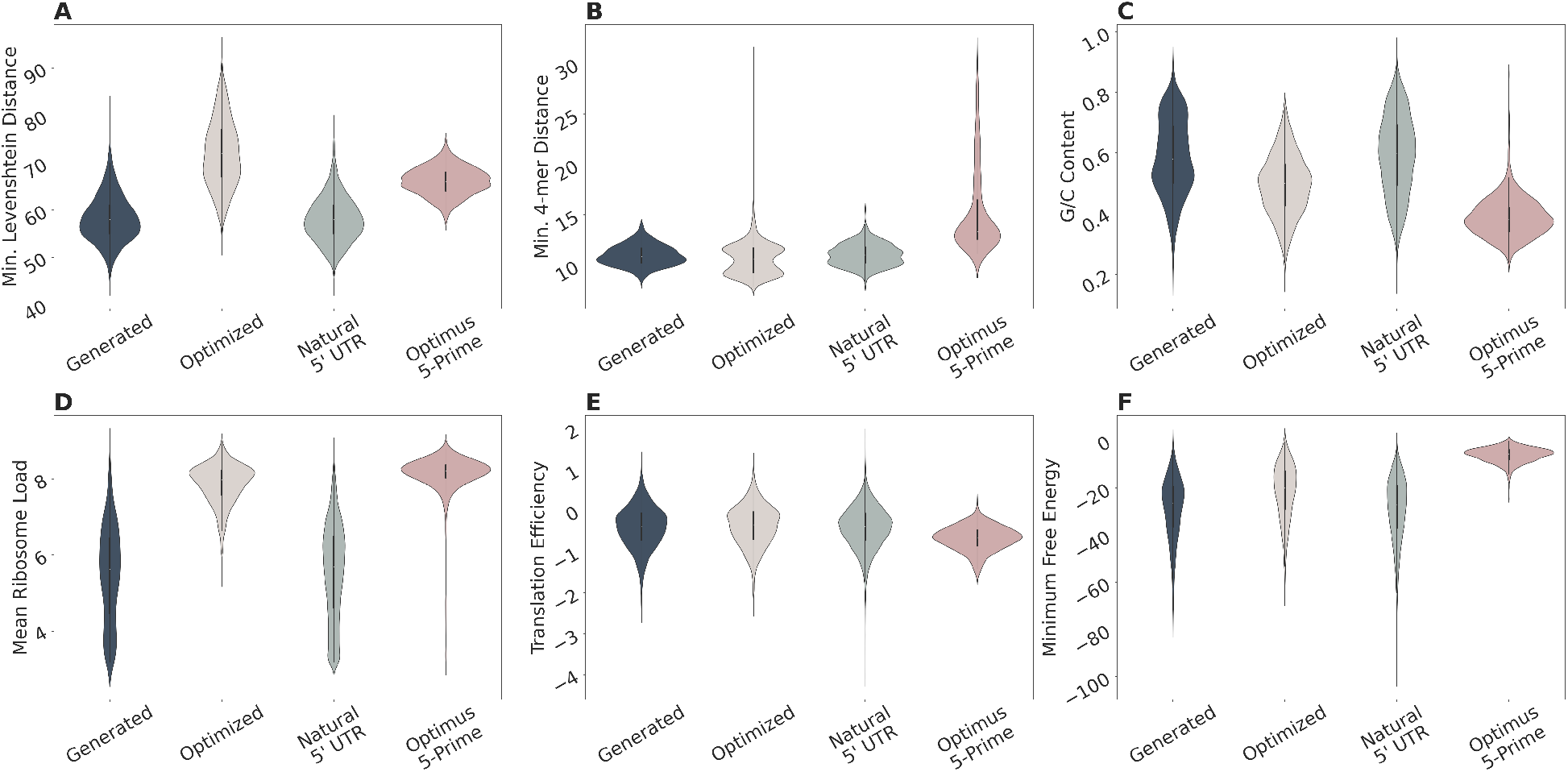
Comparison of UTRGAN-generated, natural, UTRGAN-optimized, and Optimus 5Prime-optimized sequences. **A**) The distributions of Levenshtein distances to the closest non-identical natural samples are shown for each group (UTRGAN-generated vs. natural: *p* > 0.01, effect size = 0.01, confidence interval = (0, 0); UTRGAN-optimized vs. natural: *p* = 0.0, effect size = 0.84, confidence interval = (8, 9); Optimus 5-Prime-optimized vs. natural: p = 0.0, effect size = 0.9, confidence interval = (13, 14)). **B**) The distributions of the 4-mer frequency distances to the closest non-identical natural sample are shown for each group (UTRGAN-generated vs. natural: p = 0.51, effect size = −0.02, confidence interval = (−0.18, 0.09); UTRGAN-optimized vs. natural: p < 0.01, effect size = −0.18, confidence interval = (−0.61, −0.31); Optimus 5-Prime-optimized vs. natural: p < 0.01, effect size = 0.89, confidence interval = (2.53, 2.92)). **C**) The GC content distributions are shown for each group (UTRGAN-generated vs. natural: p = 0.12, effect size = −0.02, confidence interval = (−0.01, 0.008); UTRGAN-optimized vs. natural: p < 0.01, effect size = −0.42, confidence interval = (−0.10, −0.09); Optimus 5-Prime-optimized vs. natural: p = 0.0, effect size = −0.81, confidence interval = (−0.21, −0.20)). **D**) The predicted MRL distributions are shown for each group (UTRGAN-generated vs. natural: p = 0.09, effect size = −0.03, confidence interval = (−0.14, −0.01); UTRGAN-optimized vs. natural: p = 0.0, effect size = 0.92, confidence interval = (2.14, 2.27); Optimus 5-Prime-optimized vs. natural: p = 0.0, effect size = 0.92, confidence interval = (2.37, 2.5)). **E**) The predicted translation efficiency distributions are shown for each group (UTRGAN-generated vs. natural: p = 0.92, effect size = 0.001, confidence interval = (−0.01, 0.01); UTRGAN-optimized vs. natural: p = 0.011, effect size = 0.04, confidence interval = (−0.61, −0.31); Optimus 5-Prime-optimized vs. natural: p < 0.01, effect size = −0.37, confidence interval = (0.006, 0.053)). **F**) The distributions of predicted MFEs are shown for each group (UTRGAN-generated vs. natural: p = 0.42, effect size = 0.01, confidence interval = (−0.5, 1.1); UTRGAN-optimized vs. natural: p < 0.01, effect size = 0.32, confidence interval = (6.29, 7.8); Optimus 5-Prime-optimized vs. natural: p = 0.0, effect size = 0.96, confidence interval = (20.69, 21.99)).

### 2.3 The generative model maintains the 4-mer distribution of the sequences

Another metric used to measure the similarity among two sets of sequences is the distribution of the k-mers within the sequences. We compute the frequency of the 4-mers for each sequence as also done in the literature [64]. Then, for each sequence in the natural and generated sequences, and the sequences optimized by UTRGAN and Optimus 5-Prime, we find the closest sequence (Euclidean distance) in the entire set of natural 5’ UTRs, ignoring identical sequences (so natural sequences match the nearest neighbor). We obtain a distribution per group. We discard a few of the generated sequences due to their anomalously lower distances to the set of natural sequences.

We observe very similar distance distributions for the generated and natural samples as shown in Figure 2B, indicating that the generated set of sequences retain structural similarity to natural sequences and yet, are distinct in terms of the sequence, as also shown in the previous subsection. In accordance, we observe that the sequences optimized by Optimus 5-Prime have a different 4-mer distribution despite being similar to natural sequences in terms of the sequence itself.

### 2.4 GC content distribution of the generated 5’ UTRs resemble natural 5’ UTRs

The GC content of a sequence is an important aspect for the stability molecule [35]. We observe that the mean GC contents in the generated and natural samples are very similar, and the generated sequences are diverse with respect to their GC contents (Figure 2C). The UTRGAN-optimized sequences follow a similar distribution to that of natural sequences but have a lower GC content mean, indicating a link between GC content stability and MRL. This is in contrast to the GC content of the sequences optimized by Optimus 5-Prime, which follow a substantially different distribution.

### 2.5 Predicted Mean Ribosome Load and translation efficiency for the natural and generated sequences have similar distributions

While the above-mentioned metrics are informative about the performance of the model in generating samples structurally similar to the natural sequences, they do not reflect information on the function. MRL is a metric defined based on the ribosome count associated with an mRNA molecule and is considered a proxy for translation rate [33]. We use MRL to measure the similarity of the sequences from a functional perspective.

To estimate the MRL of a given 5’ UTR sequence, we use a convolutional neural network model (FramePool) that predicts the MRL of a 5’ UTR sequence [33]. Figure 2D shows that the MRL distributions of the natural and the generated sequences are almost identical. We also show the predicted MRL distribution of the UTRGANoptimized and Optimus 5-Prime-optimized sequences. As expected, both sets of sequences have a very high average MRL as they both use the FramePool [33] model to guide the optimization. However, Optimus 5Prime fails to improve the expected MRL for all sequences, which is reflected by the very long low tail in the distribution. This method introduces random mutations in the UTR sequence, and unlike our gradient-based optimization, this may degrade the MRL for many sequences.

Similarly, we use the MTtrans model trained on ribosome profiling datasets to predict the translation efficiency of the UTR sequences [76]. Translation efficiency is another proxy for the rate at which a cell translates mRNA. MTtrans is trained on various translation profiling datasets, including (i) 3 massively parallel assay (MPRA) polysome profiling datasets and (ii) 3 ribosome profiling datasets. The model trained on MPRA datasets is called *3M*, the model trained on ribosome profiling datasets is called *3R*, and the model trained on all datasets is called *3M3R*. We use the MTtrans 3R version. As shown in Figure 2E, the predicted TE for generated and UTRGAN-optimized sequences are in the same range as the natural sequences. The sequences optimized by Optimus 5-Prime, on the other hand, have a lower average predicted TE and are not distributed similarly compared to the natural sequences.

### 2.6 Generated and natural samples have similar Minimum Free Energy distributions

The MFE of a 5’ UTR sequence as part of the RNA sequence is another informative characteristic of the sequence regarding its stability. It depends on the number, composition, and arrangement of the nucleotides in the mRNA sequence [62] and the 5’ UTR in our case. Similar to the GC content, even though MFE might not be the sole indicator of any characteristic of a given 5’ UTR sequence, we expect to see similar distributions of MFE values in a large number of samples. We calculate the MFE of the sequences we use two packages Nupack [21] and ViennaRNA [31, 42]. The result with ViennaRNA package is presented as the results are similar.

As shown in Figure 2F, the distribution of this value is very similar in generated and natural sequences while very different in the set of the sequences optimized by Optimus 5-Prime. While the UTRGAN-optimized sequences are not exactly similar to natural sequences in the distribution of their MFE, these sequences cover a much wider range of MFE values compared to sequences optimized by Optimus 5-Prime.

### 2.7 Optimization yields sequences with higher expected expression

The optimization procedure, as explained in Section 4, is based on an iterative procedure that updates the input noise of the GAN to generate 5’ UTR sequences with higher expression. We use the model to fine-tune the sequences for higher predicted expression. The model used for prediction is a convolutional neural networkbased model (Xpresso [1]) that outputs the predicted log TPM expression for the input DNA sequence. The mean saliency scores in the Xpresso paper [1] are used to measure which nucleotides around the TSS affect the predicted expression more. The authors show that the most important ones are downstream of the TSS, where the 5’ UTR is placed. Based on this analysis of the Xpresso model, we expect the optimization of the 5’ UTR part of the DNA alone to increase the expression value.

As shown in Figure 3A, the optimization of 100 generated 5’ UTR sequences for maximizing the average expression of a set of 8 randomly selected Human genes (*MYOC, TIGD4, ATP6V1B2, TAGLN, COX7A2L, IFNGR2, TNFRSF21, SETD6*) results in increased average predicted mRNA expression in 80% of the DNA samples with the optimized 5’ UTRs. To calculate a single score for each 5’ UTR, we average the expression over all 8 genes with the generated 5’ UTRs replacing the original ones. Note that we are limited by updating the 5’ UTR only, and it is not the sole determinant of the expression. Yet, we observe up to 61% increase in the average predicted gene expression compared to the initial value. In 20% of the cases, the optimization results in reduced expression compared to the initial generated sequence. In those cases, the user can pick the initially generated 5’ UTR. We discuss the possible reasons for degradation in the Discussion Section. In addition, our optimized 5’ UTR resulted in a 38% increase in the average predicted expression of a distinct set of 8 randomly selected genes (*ANTXR2, NFIL3, UNC13D, DHRS2, RPS13, HBD, METAP1D, NCALD*). This indicates that the optimization is capable of generalizing 5’ UTRs for higher average gene expression.

**Fig. 3:**
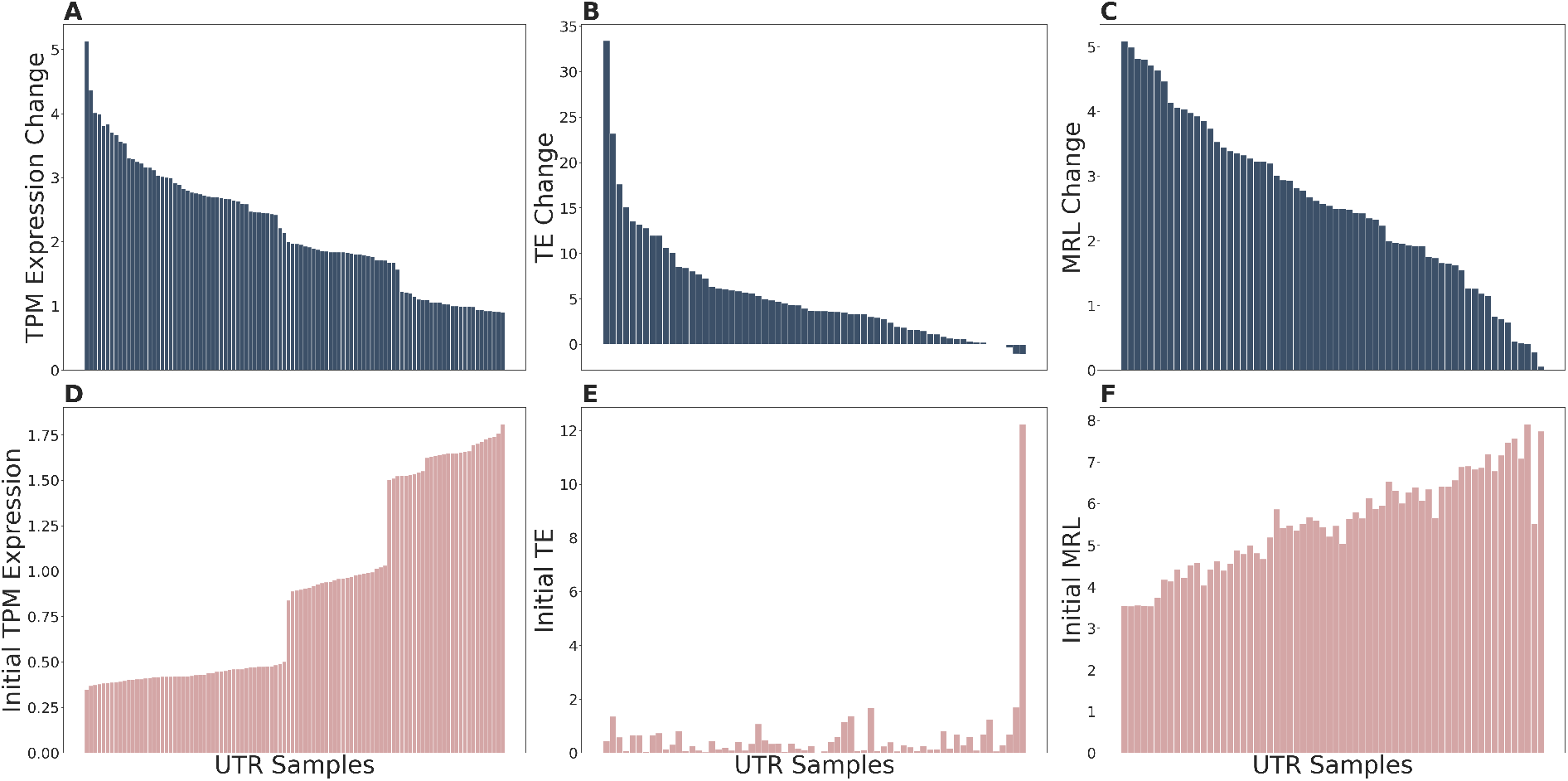
Overall performance of gene expression, TE, and MRL optimization. **A)** The expression change after 3,000 iterations of optimization for 8 DNA samples shows that the model successfully generates 5’ UTR sequences and optimizes a majority of those to improve the original expression. The values are TPM Expression values, and the optimization yields higher predicted gene expression in 80% of the DNAs. **B)** Here is shown the translation efficiency change of 64 optimized generated 5’ UTR sequences. In this case, a 97% majority of the sequences yield substantially higher TE after optimization. **C)** Similar to gene expression and TE, optimization is effective for the majority of the sequences and performs better for sequences with low initial MRL values. **D, E, and F)** Here, we show the initial gene expression, TE, and MRL values for the corresponding samples in panels D, E, and F, respectively. For all three panels, initial values tend to increase towards the right. We see that it is more likely for the optimization to degrade performance for samples with high initial TE or gene expression values. Unlike gene expression and TE, degradation in the predicted MRL of the optimized 5’ UTRs is rare.

### 2.8 Mean Ribosome Load and Translation Efficiency increases after optimization

Unlike the mRNA abundance, the MRL is not a DNA-specific metric and does not require a target DNA or mRNA sequence for optimization. The FramePool [33] model can predict the MRL value for 5’ UTR sequences of any length and does not require the UTR sequence to be attached to an mRNA sequence. We use this model and optimize the initially generated 64 5’ UTRs for 10,000 iterations. The results show increased MRL in more than 95% of the generated 5’ UTRs (See Figure 4A). The number of used sequences can be increased, but we would expect similar results as each sequence is optimized separately, and the gradients are not merged. We also demonstrate in Supplementary Figure 2, that the model learns a meaningful latent space with respect to the predicted MRL values for the generated and optimized sequences.

**Fig. 4:**
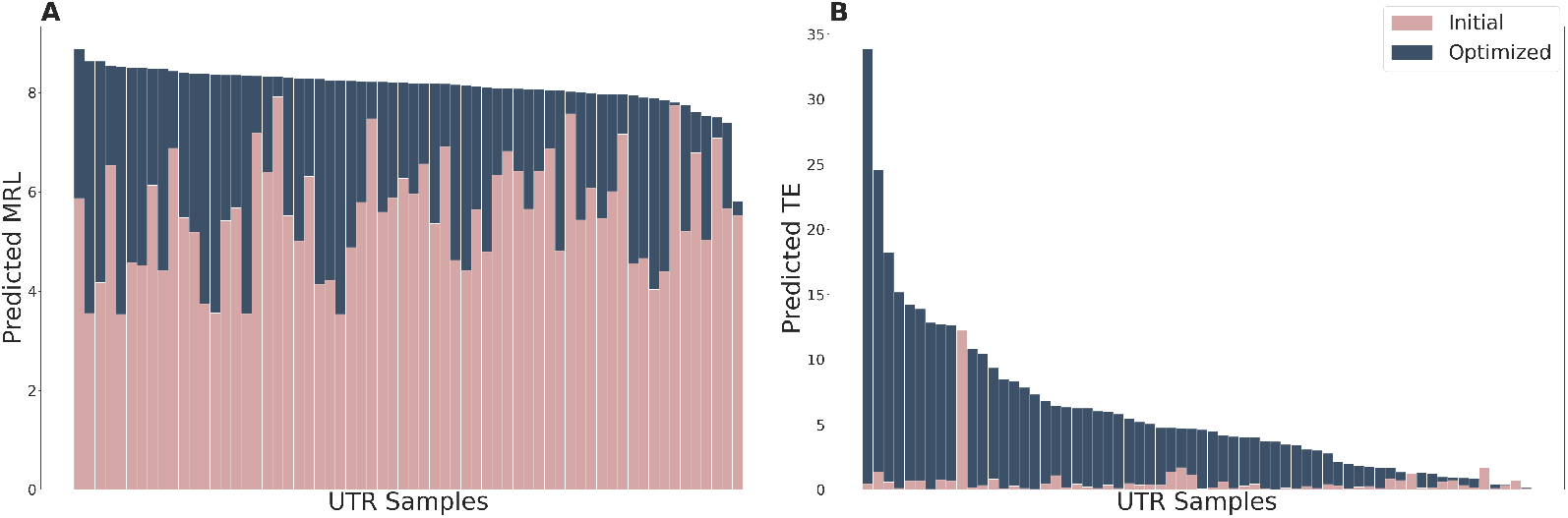
Overall performance of MRL and TE optimization. **A)**. The optimization for higher MRL using FramePool results in sequences with a considerably high average MRL of 7.6. The optimization result may vary depending on the initial random sequences, and it optimizes almost all sequences to MRLs higher than 7. **B)**. Translation efficiency reaches a high value of 33 after optimization. The initial values are all around the average TE of the natural samples, and optimization increases the average more than 32-fold, and the highest optimized value is close to the maximum value of the natural sequences. The optimized values here are behind the initial values in all panels, and for the few sequences where the blue bar is not shown, the optimized value is smaller than the initial value.

Another model we use to optimize our generated 5’ UTRs is the MTtrans model [76], which predicts translation efficiency. Translation efficiency is another proxy for the rate that a cell translates mRNA. MTtrans is trained on various translation profiling datasets, including (i) 3 massively parallel assay (MPRA) polysome profiling datasets and (ii) 3 ribosome profiling datasets. The model trained on MPRA datasets is called *3M*, the model trained on ribosome profiling datasets is called *3R*, and the model trained on all datasets is called *3M3R*. Here, we use the MTtrans *3R* model to optimize generated 5’ UTRs for higher translation efficiency (TE). The results in Figure 4B show that the optimization yields a substantial increase in TE values. To combine the optimization for mRNA abundance and translation efficiency, one can optimize many sequences for gene expression first and then select the ones also with higher MRL or translation efficiency. We also present our two-step optimization results in Section 2.11.

In addition to the TE prediction using the 3R model, the MTtrans 3M model allows us to extract important motives from the MPRA datasets used for training the model [57]. It outputs a set of 256 7-mers as motives that affect the predicted MRL more than others, both positively and negatively. During MRL optimization using the FramePool model, we observed that the number of motives affecting the MRL negatively was reduced by 53% on average after optimization, while positive motives were mostly preserved or increased. This shows that optimization using our model eliminates the negative motives based on the gradients backpropagated from the MRL prediction model.

To further analyze the results of the optimization, we compare different characteristics of the generated sequences and the optimized ones for both TE (See Supplementary Figure 3) and MRL (See Supplementary Figure 4) optimizations. We see that both TE and MRL optimization favor longer UTR sequences and lower GC content. In terms of MFE, both optimizations reduce the average absolute MFE of the sequences slightly. However, there is no clear correlation between the predicted TE and MFE values in both cases.

### 2.9 Optimization improves the expected expression for specific target genes

As discussed above, our optimization procedure increases the gene expression when we optimize multiple generated 5’ UTRs for multiple gene sequences (optimization based on average). Instead, we can work with a single target gene as well. To optimize 5’ UTRs for a specific gene, we generate 64 sequences using a fixed seed and optimize them for higher expressions when placing the UTR of the target gene only.

We optimize 5’ UTRs for the *TLR6, IFNG, TNF*, and the *TP53* genes from the human genome. Noreen and Arshad [48] discuss the role of the *TLR6* gene in the regulation of innate as well as adaptive immunity. Based on its role in the immune system, the expression level of this gene can affect the immune response. *IFNG* (*IFN-γ*) gene also has a role in the immune system and encodes an important cytokine, interferon-gamma, for immune response. [9, 61]. Similarly, *TNF* or *TNF-α* is the tumor necrosis factor gene and encodes cytokines that play a crucial role in the immune system’s inflammatory response. Finally, *TP53* gene is one of the most important tumor suppressor genes and encodes the P53 protein [37, 49]. Controlling the expression of these genes has important implications in the immunology and oncology fields. These are provided as examples to show the capacity of the model to increase the expression of specific target genes. This approach can be used on any gene for expression optimization, both to increase and decrease expression.

We show in Figure 5 that optimization for these genes improves gene expression for more than 95% of the generated 5’ UTRs. It provides a 4-fold, 37%, 4.2-fold, and 63% increase in expression on average for each gene, respectively. These results show that optimization is successful on average, and users can pick the top generated sequence for their application. The performance increase for the highest expression yielding 5’ UTR is 8-fold, 3-fold, 32-fold, and 3.3-fold for each respective gene. This optimization results in 2.2-fold increase in the average expression of the mentioned 4 genes together. The sequences of the best-performing 5’ UTRs for these genes are provided in Supplementary Table 1.

**Fig. 5:**
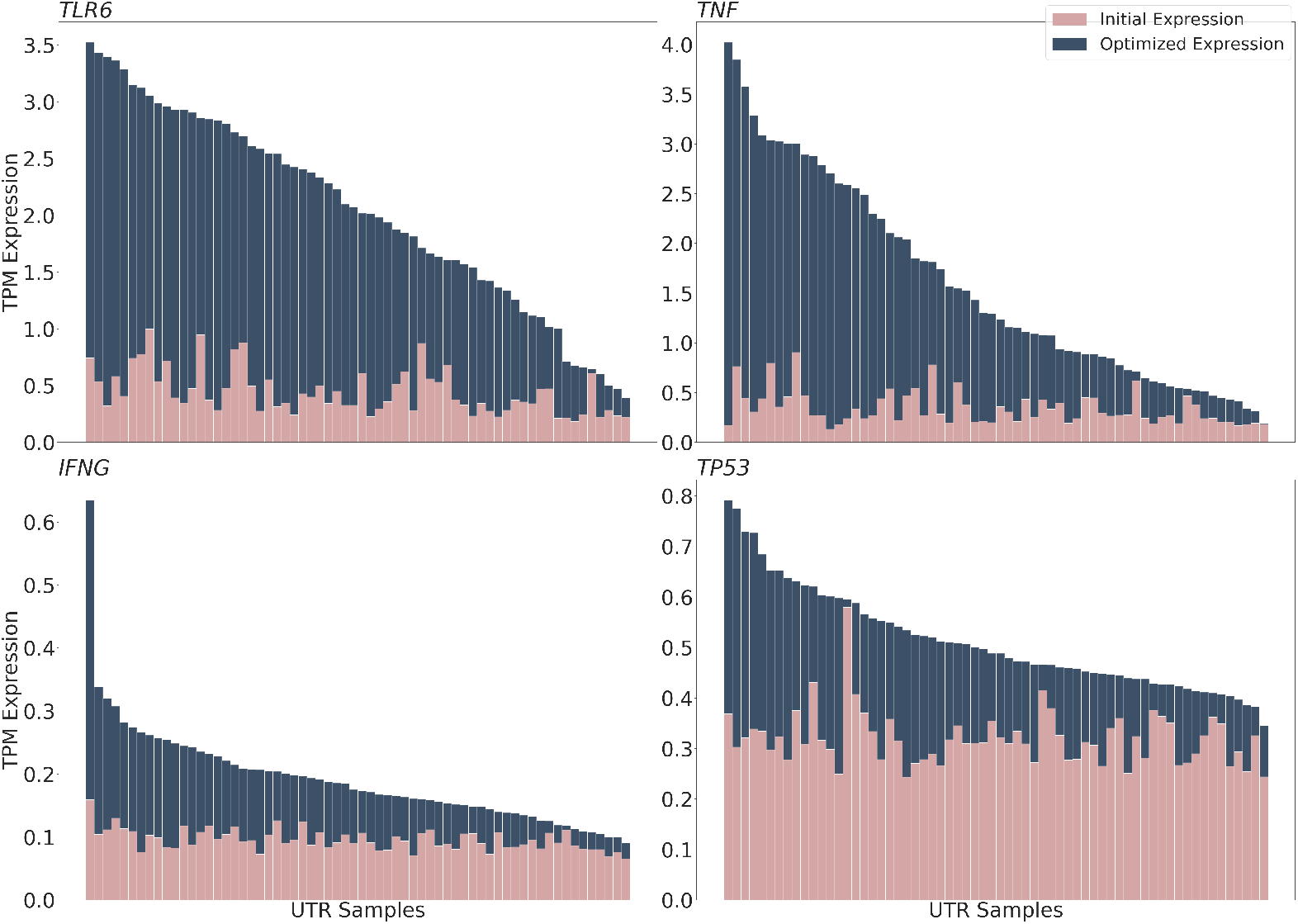
Expression optimization for specific target genes. In addition to optimizing for random genes, it is possible to optimize many generated UTR sequences for specific genes. Here is shown the result of optimization for four specific genes. *TLR6, IFNG, TNF*, and *TP53* genes show a substantial increase in the expression for optimized sequences compared to initial predicted expression values. The optimization shows 4-fold, 37%, 63%, and 4.2-fold increase in expression on average for *TLR6, INFG, TNF*, and *TP53*, respectively. The optimized expression values are behind the initial values in all panels, and for the few sequences where the blue bar is not shown, the optimized value is smaller than the initial value.

Moreover, we observe that the GC content of the sequences increases with gene expression optimization, and the best-performing sequences exhibit GC content as high as 85 percent. The GC content of some natural sequences even exceeds this number. It should be noted that even though high GC content can lead to higher mRNA expression levels [3] [36], it may not be desirable in all circumstances due to various concerns such as stability [35]. To address this, we incorporate a control mechanism in our optimization process, allowing us to limit the maximum level of GC content for the selected best-performing sequences. For optimization results with an upper bound of 65 percent for the GC content of the optimized sequences, please refer to Supplementary Figure 1. The procedure still substantially increases the expected expression of all four genes.

### 2.10 Synthetic 5’ UTRs yield higher predicted expression for target genes compared to their natural 5’ UTRs

Our main goal in this study is not to replace existing 5’ UTRs of natural genes. The experiments in the previous subsections with a single or a set of target genes are to show that the optimization mechanism can tailor the initially generated (synthetic) 5’ UTR sequences toward a goal (maximize expression) using natural gene sequences as templates/examples when their UTR sequences are replaced with the generated and the optimized UTR sequences. Our approach is rather intended for novel and synthetic DNA molecules with some desired properties, such as binding to another protein after translation.

Yet, in this subsection, we investigate if the 5’ UTRs we generate yield higher predicted expression compared to the natural 5’ UTRs. For each of the *TLR6, IFNG, TNF*, and *TP53* genes, we generate 64 5’ UTR sequences and pick the sequence that maximizes the expected expression and compare this value with the expected expression of the gene using their natural 5’ UTRs. We observe that the sequences we generate yield 8-fold, 3-fold, 32-fold, and 3.3-fold increases in the expected expression, respectively.

### 2.11 Joint optimization results in both higher translation efficiency and gene expression

As shown in Sections 2.7 and 2.8, independent optimizations for translation efficiency and mRNA abundance yield positive results. Nevertheless, optimizing one does not necessarily increase the other. Here, we also consider the case where optimizing 5’ UTR sequences for both higher translation rate and mRNA abundance for a target gene. We use the same set of 4 genes used in Section 2.9. We use a sequential optimization procedure to achieve this goal. Since TE optimization is gene agnostic and mRNA abundance optimization is gene-specific, we first optimize the sequences *z* for higher translation efficiency for 1,000 iterations. Then, we use the optimized *z* as the starting point for mRNA abundance optimization. The optimization first increases the average TE from negative to positive. That is, the average value is increased from 0.27 to 30.9. Then, the TE-optimized sequence is further optimized for mRNA abundance for each of the target genes for 1,000 iterations. Although the second optimization procedure slightly decreases the average translation efficiency from 1.49 to slightly above zero for certain genes, both the translation efficiency and mRNA abundance are higher compared to the initially generated sequences, as shown in Figure 6. In addition, we observe that our two-step optimization performs better than optimizing for these two scores simultaneously. The sequences of the best-performing 5’ UTRs after joint optimization for these genes are provided in Supplementary Table 2.

**Fig. 6:**
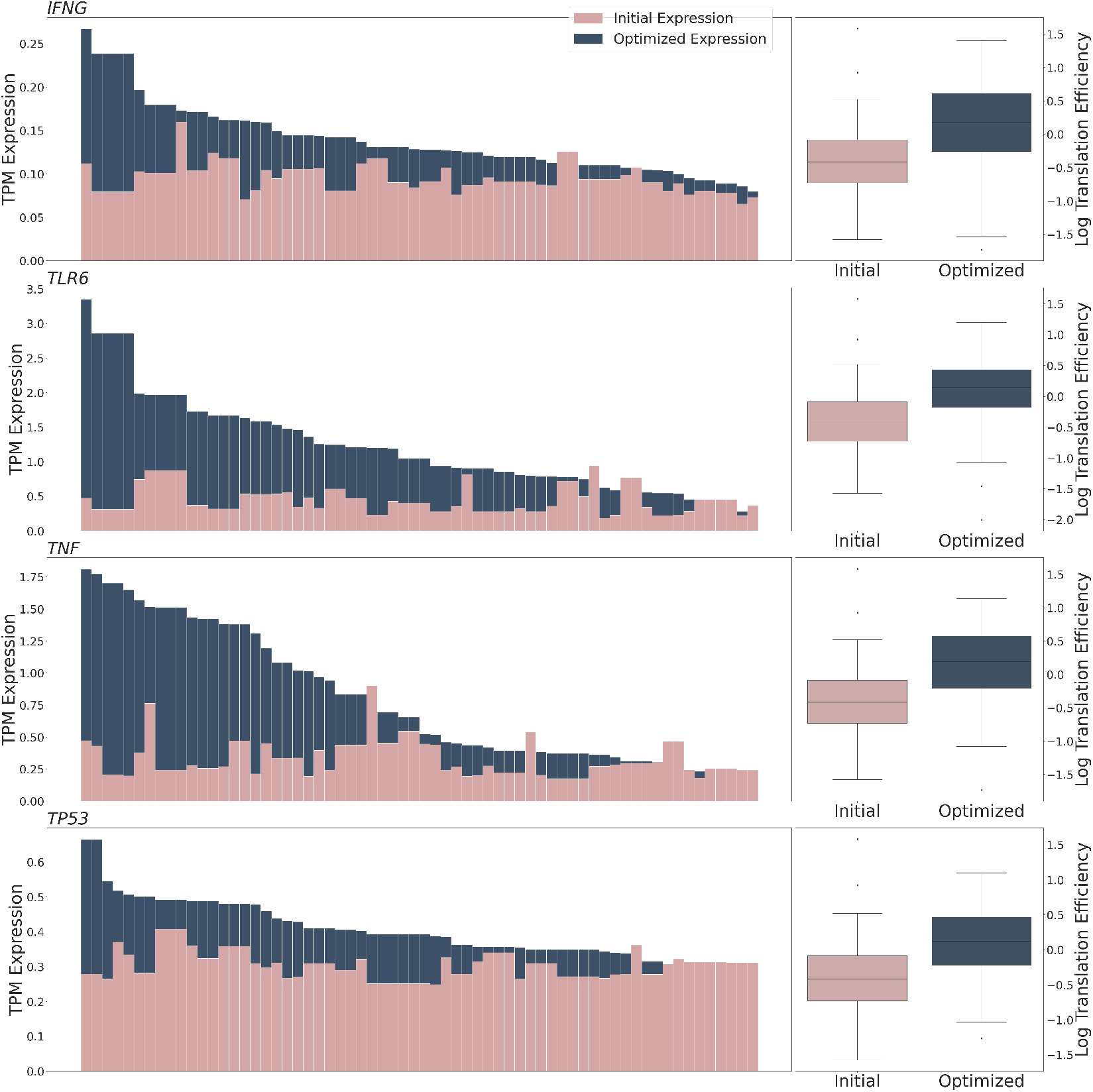
Jointly optimizing for TE and mRNA expression. The generated sequences are first optimized for high TE for 1,000 iterations, and then the sequences resulting from the optimization are used as the starting latent vector for gene expression optimization for another 1,000 iterations. As seen for the four genes, *TLR6, IFNG, TNF*, and *TP53*, the two-step optimization results in higher gene expression and translation efficiency. The predicted gene expression increases up to 12-fold, 1.8-fold, 10-fold, and 2.8-fold with respect to the natural 5’ UTRs of the genes, respectively. The maximum predicted TE after optimization is at least 77% higher than the natural 5’ UTR of the target genes. The optimized expression values are behind the initial values in all panels, and for the few sequences where the blue bar is not shown, the optimized value is smaller than the initial value. The box plot characterizes the TE values using the 25th, 50th, and 75th, also known as quartiles (Q1, median, Q3) and the interquartile range (IQR = Q3 - Q1), with whiskers extending to a maximum of 1.5 times the IQR. Outliers beyond the whiskers are plotted separately.

### 2.12 Regulatory Elements are Conserved in UTRGAN-generated and -optimized Sequences

One of the most important regulatory elements in 5’ UTRs are uORFs, which play various roles, such as translation reinitiation [7, 16, 32]. We obtain the set of known human uORF sequences from uORFdb [44] (n = 2,422,112) and search for human uORFs with as short as 7bp and as long as 64bp, which includes 402,140 sequences, in the set of sequences of interest. Figure8A shows the number of uORFs found given in each sequence set. Note that we have a larger set of natural UTRs, so the corresponding count is normalized. We observe that there are a substantial number of uORF elements present in our UTRGAN-generated/optimized sequences, while very few of them are maintained in the sequences optimized by Optimus 5-Prime.

IRES sequences are another set of RNA elements utilized in cap-independent translation as part of the protein synthesis process [59]. IRES sequences are most frequently found in 5’ UTR regions and are used by cells to increase the translation rate of certain genes [34]. Here, we obtain the human IRES elements from the IRESBase database [75] and compare the sequence sets of interest with respect to their maximum pairwise alignment scores to IRES elements. We use alignment as IRES sequences tend to be relatively long (longer than 128bp). As shown in Figure8D, UTRGAN-optimized sequences have substantially higher alignment scores compared to the generated ones and sequences optimized by Optimus 5-Prime.

We also investigate if the generated sequences are enriched with elements such as Kozak sequences and G-quadruplexes (G4) in the sequence sets of interest. G4s are elements with specific structures in G-rich regions of the 5’ UTR that affect the stability of the mRNA sequence [8,19]. We detect G4 structures using the G4Boost model [12]. Kozak sequences are usually located in the non-coding region upstream of the translation initiation site [74]. In addition to the consensus Kozak sequence (GCCGCCRCCAUGG), alternative sequences are known to play a similar regulatory role [72, 73]. However, we do not see any of the Kozak sequences in many of the natural 5’ UTRs, as shown in Figure 8C. We search for exact matches of these alternative Kozak sequences with different initiation start codons (‘AUG’, ‘AUA’, ‘CUG’, and ‘GUG’) in the sequences [46, 73]. As shown in Figure8A and Figure8B, respectively, UTRGAN-generated and optimized sequences include many instances of G-quadruplexes and Kozak sequences while in the sequences optimized using Optimus 5-Prime they rarely occur.

### 2.13 Motif Analyses Provide Insights on Learned Regulatory Patterns

We perform a motif analysis on the 1,024 UTRGAN-optimized sequences for *de-novo* motif identification. We use the MEME suite [5, 6] which yields the set of motives that are enriched given a sequence set. We retrain the top 50 motives of length 7-15 bps for each sequence set of interest (UTRGAN-generated/optimized and Optimus 5-Prime-optimized). We use the same tool to identify the top 50 important motives in natural 5’ UTRs, human uORF sequences, and IRES sequences. Then, we use the TomTom motif comparison tool [30] to find the top-3 motif matches across these two groups (e.g., UTRGAN-optimized vs IRES). In Figure 7, we show the top matches between the natural sequences (5’ UTR, uORFs, and IRES) and UTRGAN-optimized sequences (See Supplementary Figure 6 for the same comparison for UTRGAN-generated sequences). Analyzing these motives, we observe that some motives in generated sequences seem to disappear or get modified during the optimization as the *de novo* motif identification results in different motives for generated and optimized sequences. These results show that UTRGAN successfully maintains important motives found in natural sequences, and optimization results in meaningful changes in the number of present regulatory elements for high efficiency in the desired target.

**Fig. 7:**
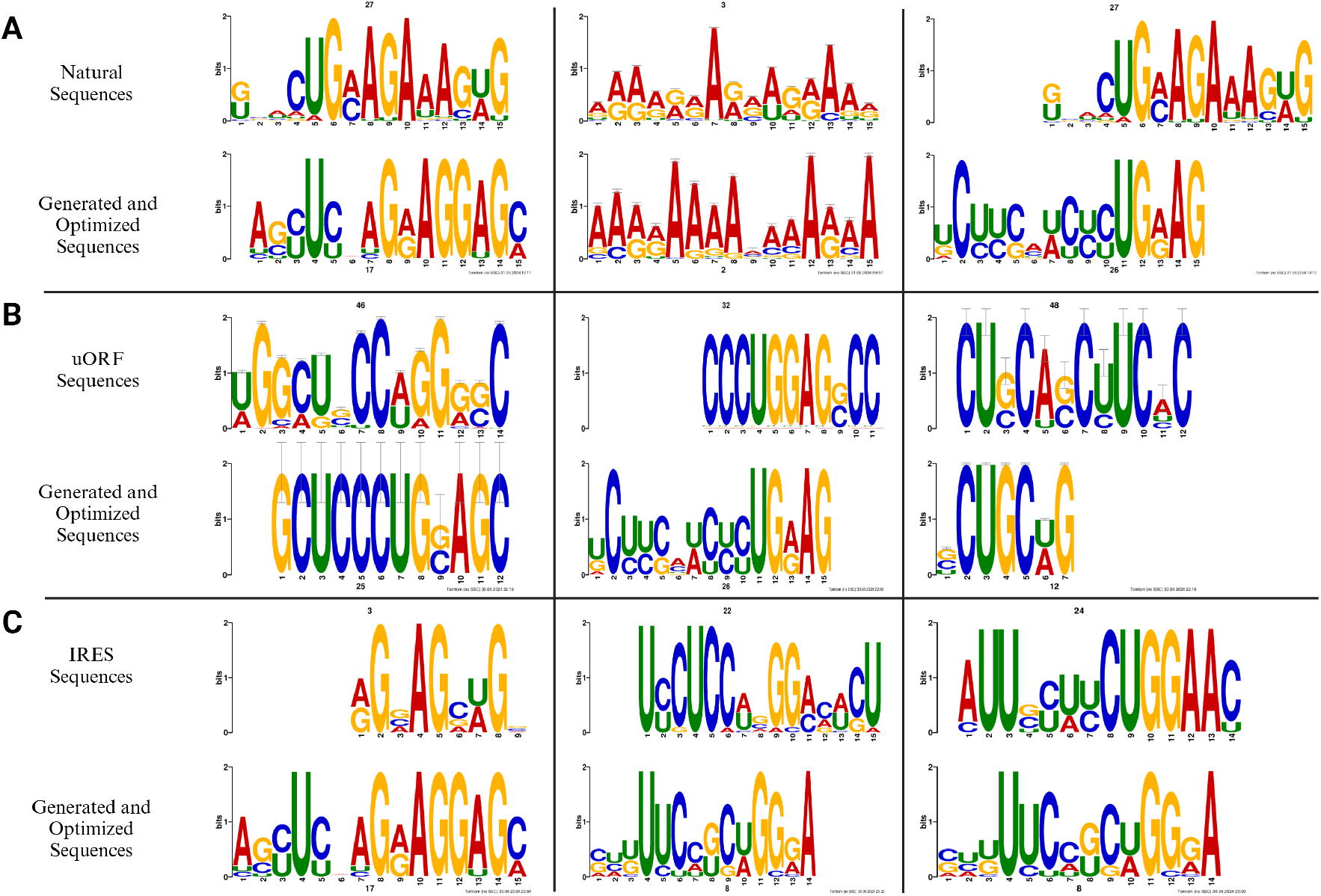
Top Matching Motives between UTRGAN-optimized and various natural sequences. The top-3 motif matches between the sequences optimized using UTRGAN and the natural 5’ UTR, uORF, and IRES sequences are shown on panels **A**,**B**, and **C**, respectively. The leftmost motif is the top match for each category.

**Fig. 8:**
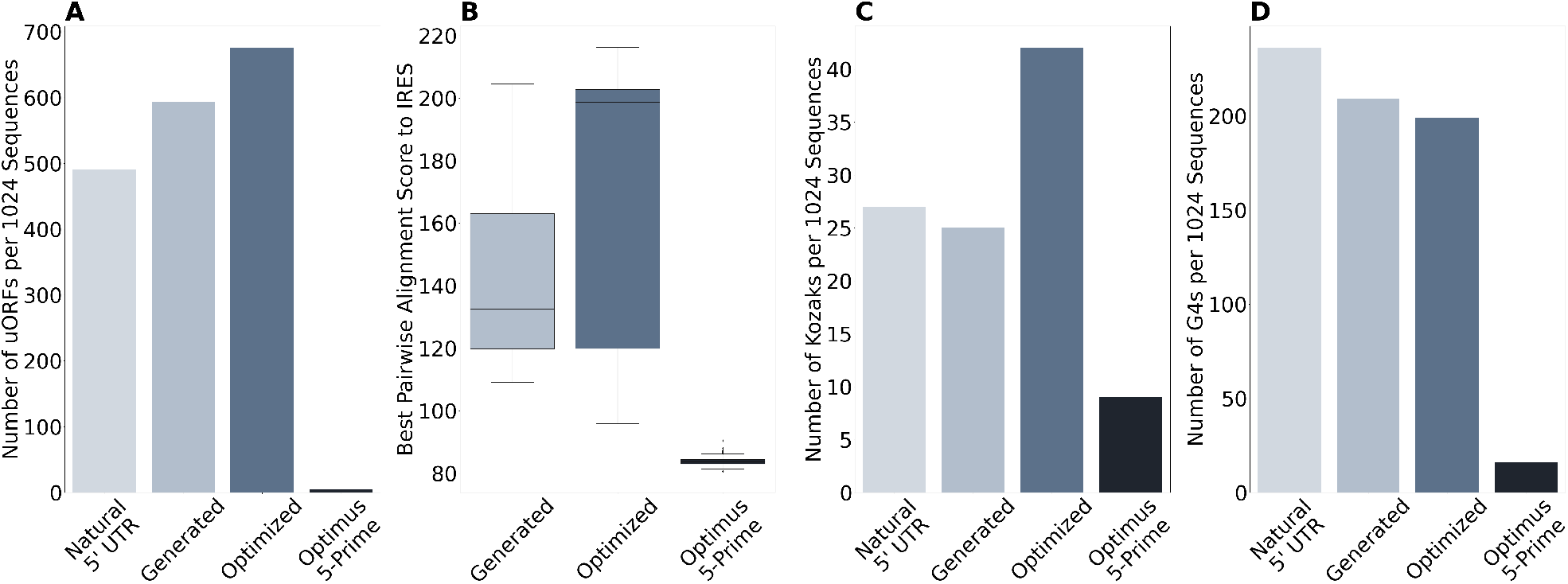
Regulatory element analysis of UTRGAN-generated, UTRGAN-optimized, Optimus 5-Prime-optimized and natural 5’ UTR sequences. The elements analyzed here include uORF, IRES, Alternative Kozak, and G4 sequences, which are shown on panels **A**,**B**,**C**, and **D**, respectively. We count the occurrence of each regulatory element in natural 5’ UTR, generated, UTRGAN-optimized, and Optimus 5-Prime-optimized sequence sets. Note that all sets except the natural 5’ UTR set have 1,024 sequences. So, we normalize the latter to show a number of hits for 1,024 sequences on average.

We also observe that UTRGAN-optimized sequences result in more ‘U’ rich motives in contrast to the initially generated sequences that are ‘A’ rich instead. Studies show that ‘U’ rich sequences in cancerous cells are overregulated, and there is a correlation between the percentage of ‘U’ and high translation in related genes [45, 71]. In line with this observation, our optimization scheme aims for patterns with higher ‘U’ content to achieve a higher translation rate.

### 2.14 The cytotoxic effect of TNF-*α* proteins containing synthetic UTRs exhibit higher translation

Via in-vitro experiments, we test the efficiency of the designed UTRs. We compare the translation rate of the TNF-*α* protein when using (i) the UTRGAN-generated/optimized 5’ UTRs, and (ii) the human *β*-globin 5’ UTR. TNF-*α* is a pleiotropic cytokine involved in the various physiopathological processes and is known to induce cytotoxicity in select target genes, leading to cell death. The MCF-7/MX cell line is known to be vulnerable to TNF-*α* protein. We quantify the effect of the synthetically generated UTRs on the translation rate of this protein by using the cytotoxic effect on MCF7 cells as a proxy.

We optimize the 5’ UTR sequences with respect to the mean ribosomal load (See Methods for details). To be sure about the initiation of translation, we manually add the consensus Kozak sequence (GCCGCCACCAUGG) at the 3’ end of the 5’ UTRs (both synthetic and natural). Consequently, we design an mRNA containing the 5’ UTR region with the full-length protein sequence, encompassing the first 76 amino acids of the TNF-*α* protein. This specific region acts as a leader sequence for the protein’s secretion in its natural process [68]. Subsequently, we transcribe these mRNAs in-vitro and use them to transfect HEK293T cells, which possess a high protein production capacity and the necessary elements for secretion, such as ADAM10 and ADAM17 sheddases [52].

We conduct all experiments under precisely controlled conditions for all mRNA samples, ensuring that any observed effects were solely attributable to the 5’ UTRs. Given that TNF-*α* induces a cytotoxic effect on MCF7 cells and inhibits their proliferation, we design an assay to quantify the TNF-*α* produced from given in-vitro transcribed mRNAs by comparing the viabilities of the MCF7 cells. According to the viability results, as shown in Figure 9, all of the UTRGAN-generated and then optimized 5’ MRL UTRs exhibit a significantly higher cytotoxic effect than the 5’ human *β*-globin UTR at a 90 percent confidence level where three of them have a 95 percent confidence level with respect to the one-tailed Student’s t-test.

**Fig. 9:**
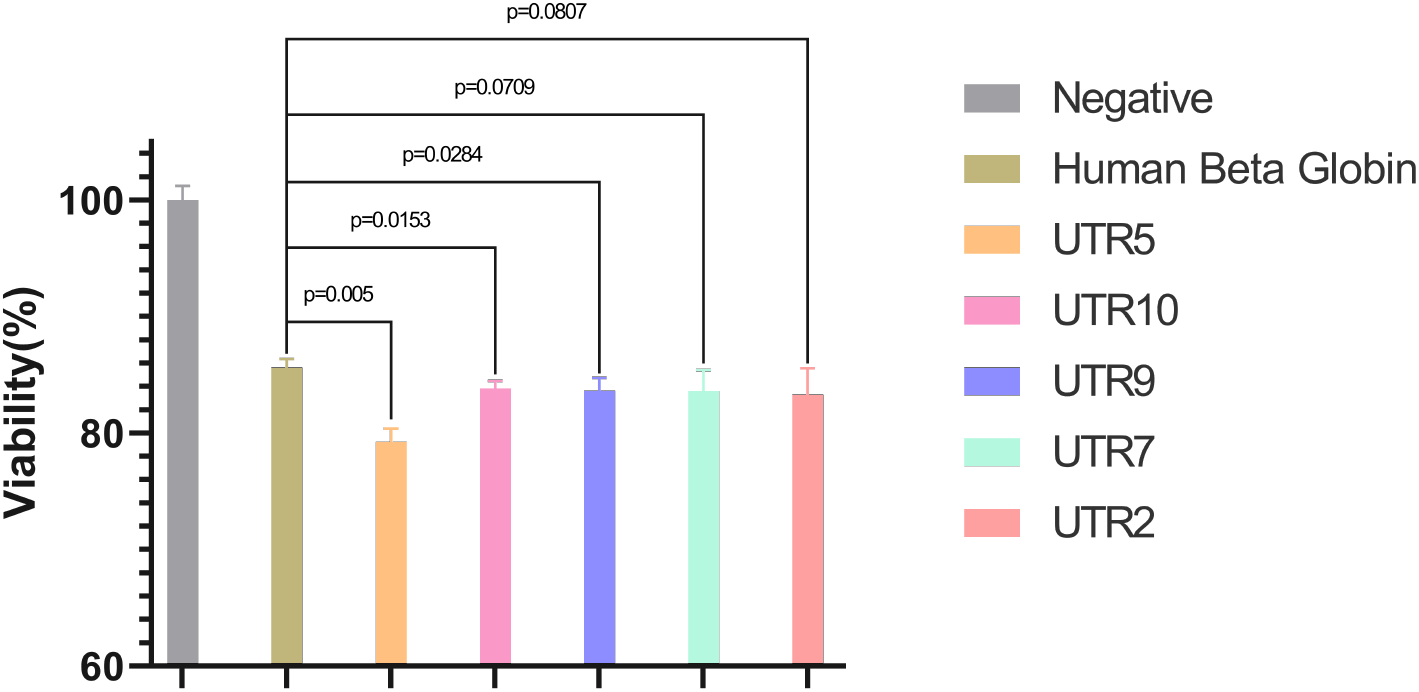
Cytotoxicity level of TNF-*α* with natural and UTRGAN-generated and then UTRGAN-optimized 5’ UTRs. Shown here are the effects of TNF-*α* proteins in transfecting MCF7 cells. These values show the level of translation using different 5’ UTRs, including the human *β*-globin and 5 generated/optimized UTRs. The designed UTRs are more effective than the human *β*-globin UTR in upregulating the translation of the TNF-*α* mRNA.

## 3 Discussion

We introduce UTRGAN, the first framework for developing synthetic 5’ UTR sequences optimized towards a target feature. Our approach enables us to generate sequences with similar attributes to the Human 5’ UTR samples without the need for exploring intractable search space of DNA sequences that have lengths of up to 128 nucleotides. We advance the state-of-the-art, which relies on optimizing existing UTRs via modifications and by generating sequences from scratch. Our model can also generate sequences of variable lengths, further increasing the possible space of generated sequences.

UTRGAN is a robust framework for synthetic 5’ UTR engineering and is applicable to numerous applications for genetic circuit design. The model’s components consist of a GAN adopted to be trained on the Human natural 5’ UTR dataset and state-of-the-art prediction models for MRL, TE, and mRNA abundance [33] [76] [1]. Our model can potentially optimize sequences for given any objective function as long as a differentiable prediction model exists to predict the target metric for a 5’ UTR sequence. We can maximize and minimize the target feature. Similarly, the model can work with classification models to generate sequences of a particular class if the prediction model is replaced with such a classifier.

Although UTRGAN can generate and optimize sequences with desirable characteristics, the optimization can lead to decreased TE or mRNA in less than 5% of the cases for TE in the worst optimization and 20% of the cases for mRNA abundance for optimizing UTRs for multiple genes, while it rarely happens for MRL optimization (Figure 3C). One possible reason for decreased expression is the limitation with regard to the predictor model (Xpresso). It operates on the entire DNA (gene) sequence, and while back-propagating the gradient to update the sequence, we can only use part of the gradient that corresponds to the 5’ UTR. Thus, changing the 5’ UTR but not the rest of the sequence might lead the model to end up at a worse point in the loss space, and changing the rest of the DNA is not desirable for the target application, which requires the rest of the DNA to remain unchanged. Furthermore, in both mRNA expression and TE optimization, we observe that the decreased scores tend to occur when the optimization procedure starts with a sequence that yields a high score, as seen in Figure 3D and E). In such a case, users can use the starting sequence and discard the optimization.

We expect to see wide use of data-driven sequences generated using deep learning methods in the near future. Generating 5’ UTRs utilizing this approach is practical and does not require massive computational resources. While we focus on 5’ UTRs, it is straightforward to generalize the framework to generate 3’ UTRs or other regulatory elements. It is also possible to use other generative models, such as variational autoencoders or diffusion models. Generating sequences with regulatory roles enables researchers to conduct experiments in a much shorter time, which can be particularly beneficial in mRNA vaccine production and mRNA-based therapeutics.

## 4 Methods

### 4.1 Datasets and the Experimental Setup

We obtain the natural UTR sequence data from UTRdb 2.0 [41], a complete database of 3’ and 5’ UTR sequences of various organisms. We use only the Human 5’ UTR samples. The length of the sequences ranges from 30 bp to 600 bp. However, this range is too wide for a CNN model since all the sequences should be padded up to the longest sequence, making padding longer than the sequence itself in most cases. Therefore, we limit the length of the training between 64 bp and 128 bp. This subset contains 33,250 natural 5’ UTR sequences. We use 90% of the samples for training and 10% of the samples for validation. We train the model with batches of size 64. We obtain the DNA sequences of Human genes used for optimization from the Ensembl Biomart tool [18].

We encode the data using one-hot encoding, and the dimension of each UTR is (128, 5), where 128 is the maximum size. and 5 is the dimension of the one-hot vector for each nucleotide. The DNA alphabet in one-hot encoded as follows: *A*:(1,0,0,0,0), *C* :(0,1,0,0,0), *G* :(0,0,1,0,0), *T* :(0,0,0,1,0), *X* :(0,0,0,0,1) where *X* is the padding character used when the sequence is shorter than 128 bp. This padding is necessary because the input of the Critic as a convolutional neural network has to be in fixed dimensions, and the sequences naturally vary in length. In addition, to generate truly random sequences, we select a random length between 64 and 128 nucleotides and then select each nucleotide randomly from the DNA alphabet. We use 2048 random, 2048 generated, and all the 33,250 natural sequences in all the analyses unless stated otherwise. See Supplementary Notes 2.1 for details about the hardware details and Supplementary Notes 2.2 for hyperparameter optimization. The code is available at https://github.com/ciceklab/UTRGAN.

### 4.2 The Model

#### Overall Architecture

The generative model used here is a variant of a Generative Adversarial Network (GAN) [27]. The input of the GAN is a random noise sample *z*, and we train the GAN to generate realistic 5’ UTR sequences. The GAN learns to map the noise to the 5’ UTR space. Then, we can optimize the model to generate specific sequences by optimizing the input noise only. The optimization performed here does not update the weights of the generator and changes the input noise *z* instead.

The sequences generated by the GAN are scored by the selected scoring models. We use three deep convolutional neural networks to score the generated sequences. The mRNA abundance scoring model [1] predicts the log TPM expression of a given gene sequence, including a 5’ UTR. Xpresso provides three models for predicting median expression among many cell types, K562 erythroleukemia cells and GM12878 lymphoblastoid cells [1]. We use the median prediction model in this study, but the optimization can be performed for specific cell types using the mentioned models. The models used to predict the ribosome load and translation efficiency of the 5’ UTR require only the 5’ UTR sequence as input [33] [76]. We use the MTtrans 3R model that predicts TE values as a proxy for translation rate by inputting only the 5’ UTR sequence [76]. All models and the generator are differentiable, which enables us to optimize the input *z* with Stochastic Gradient Ascent by updating it in multiple iterations. Sequences are fine-tuned for 3,000 iterations for optimizing expression and 10,000 for optimizing MRL and TE. iterations and the performance in each iteration is stored. The version with the top performance is picked as the final optimized version of that sequence.

We also attempted to train a VAE for designing UTRs, as VAEs also provide us with a latent space that we can use for optimization. We were initially unable to train a VAE using one-hot encoded input sequences as UTRGAN does. The model did not converge. As a second attempt, we tried to train a VAE using byte-pair encoding (BPE) [23] tokenized UTR sequences, but the model failed to learn the pattern of the paddings and was, in most cases, incapable of generating sequences with padding only at the end of the sequence. Our best attempts in training a VAE resulted in a model generating semantically meaningless or very biased GC content (mostly around 100 percent GC content) sequences.

#### Generative Model Architecture

The GAN used here is based on the original Wasserstein GAN (WGAN) [4] model that is a Convolutional GAN model [53] with the Wasserstein loss [22] and Gradient-Penalty that improves the stability of the training [29].

The layers of the deep learning model are modified to fit this task based on the dimensions of the data, with 128 nucleotides, each represented by a vector of length 5. The generator *G* consists of a dense layer to increase the dimension of the input noise, following five residual convolution blocks, all using the same number of channels. Finally, the output of the last residual block is fed to a convolution layer with five output channels following a softmax layer, the output of which is the one-hot encoded sequence. See Supplementary Table 6 for details on the layer dimension of the generator. The input is a random vector of length 40, sampled from a normal distribution with a mean of zero and a standard deviation of 1.0. Technically called critic, the Discriminator *D* is a convolutional neural network with five residual blocks following a dense layer with one output. See Supplementary Table 7 for details of the critic layers. Unlike the original DCGAN model [53], there is no activation function at the output of the dense layer of the critic, as the output is an unbounded score and not a probability [4]. The parameters of the critic are updated 5 times for every update performed on the generator. This means that the critic is updated 5 times more than the generator.

The initial weights of the network are sampled from a normal distribution with mean = 0 and standard deviation = 0.1. In the loss function, as shown in Equation 1, the generator learns to minimize the distance between the distribution of the critic’s scores for the natural and generated sequences. The first part of the loss function belongs to the original WGAN model [4], and the second part includes the gradient penalty and its coefficient *λ*. Here, *x* and 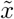 are the natural and generated samples, where we sample 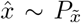 uniformly along straight lines between pairs of points sampled from the data distribution ℙ*r* and the generator distribution ℙ*g* for the computation of the gradient penalty. The gradient penalty improves the training of the GAN by enforcing the Lipschitz constraint as the model proposes [29], and this allows training for a higher number of iterations. Here, we train our model for 3,000 epochs.

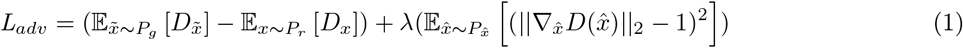

#### Expression Prediction Optimization

The Xpresso model can predict the expression of a one-hot encoded DNA sequence with a fixed length of 10,500 nucleotides. The model receives sequences of 3,500 nucleotides downstream and 7,000 nucleotides upstream of a Transcription Start Site (TSS) and predicts the log TPM expression for the sequences.

Let *z*_0_ denote the initial latent vector, then the generator inputs the latent vector (*G*(*z*_0_)) and generates the 5’ UTR, which we call *U*_*g*_. *S* denotes the gene sequence for which we design the 5’ UTR with the original 5’ UTR *U*_*r*_. The expression prediction model is denoted as *E*(*S*) where *S* is the input sequence.

To carry out the optimization, the initial 5’ UTRs are first generated using a fixed seed for the random latent vector. They are attached to the selected gene sequences, replacing the original 5’ UTRs, and the initial expression is measured. We define *R* as the function to replace the original 5’ UTR *U*_*r*_ with the generated 5’ UTR *U*_*G*_ while maintaining the length of *S* and the TSS position on the sequence (see Equation 2). The original position of the TSS in the DNA samples is the 7000th nucleotide from the start. The length of a sequence is obtained via the *len* function. We use the bracket notation to slice the sequences.

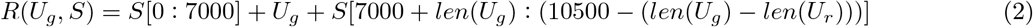

To perform the Gradient Ascent to the input latent vector for optimization, we first calculate the gradient of the output of the Xpresso model with respect to its input DNA sequences (Equation 3) and slice the part of the gradient that corresponds to the 5’ UTR from the 7000th to up to 7128th index where the 5’ UTR is located depending on *L* which is the length of the generated 5’ UTR. In the next step, the gradient of the generator is calculated with respect to the input noise vector.

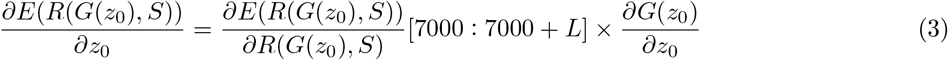

Updating the noise vector according to the gradient is supposed to increase the expression in the following iteration. This iteration is repeated 3,000 times by updating the input latent vector 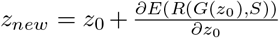. We store the history of predicted expression values for each iteration and pick the best-performing sequence as the optimized version. Although the optimization is performed on the batch of sequences, the final sequences are selected independently as if the same optimization was performed separately for each element of the latent vector. This is to enable batch optimization instead of performing optimizations on individual UTRs separately.

#### Mean Ribosome Load and Translation Efficiency Prediction Optimization

The FramePool model was trained on UTR sequences of lengths up to 100 bp. It utilizes the Frame-slice layer, which allows it to predict the MRL value for 5’ UTR sequences of any length [33]. Frame-slicing slices the sequence into 3-mers or codons, referred to as frames. It applies global pooling on the indices, which lets it work with sequences of any size. To perform the optimization with this model, again, we use the Gradient Ascent algorithm. Here, slicing is not required as the model only inputs the UTR, and the derivatives are taken directly on the 5’ UTR sequence. Considering the MRL prediction model as *M* and the initial latent vector as *z*_0_ again, the gradient is calculated as shown in Equation 4.

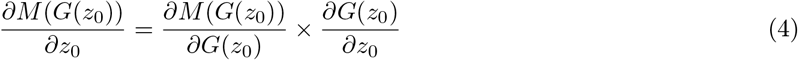

We apply the gradient by updating the input vector 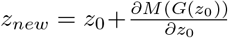. This is repeated for 10,000 iterations, and then the optimized sequences are selected as done for expression maximization and explained in Section 4.2. TE optimization using MTtrans *3R* is performed similarly with a maximum of 10,000 iterations, and the best-performing sequences are selected as mentioned above.

### 4.3 *In vitro* Experiments

As discussed in the results, we compare the generated 5’ UTRs and the human *β*-globin 5’ UTR in the translation of the TNF-*α* protein. The methodology of the conducted experiment is explained in detail in the sections below.

#### Assembling of the constructs

For the in vitro transcription (IVT) constructs, we utilized the pUC57-T7 promoter-Human -globin 5’ UTR-sfGFPHuman -globin 3’ UTR-118 poly(A) construct previously obtained from Genewiz for our laboratory. Five UTRs were selected randomly from a pool of top ten *de novo* UTRGAN-generated and UTRGAN-optimized UTRs based on mean ribosomal load (MRL). This pool of UTRs was designated as UTR1 to UTR10 (see Supplementary Table 4) to simplify labeling. The selected ones UTR2, UTR5, UTR7, UTR9, UTR10, and the human *β*-globin 5’ UTR (used as a control) were cloned into the pUC57 vector as described earlier, which includes a coding region for human TNF−*α*. The human TNF−*α* coding region was sourced from the pLI−TNF plasmid [52] (deposited to Addgene by Veit Hornung, Addgene #171179). The selected 5’ UTR regions were incorporated into the coding region through polymerase chain reaction (PCR) and then ligated into the digested pUC57 vector instead of sfGFP. Following cloning, sequence verification of the constructs was conducted using Sanger sequencing (Genewiz). Throughout the cloning process, the DH5-*α* strain of *Escherichia coli* was employed.

#### *In vitro* transcription and removal of the inorganic contaminants

The cloned plasmids were isolated using GeneJET Plasmid midiprep kit (Thermo Scientific) from overnight-grown bacterial cultures. Linearization was achieved using a single restriction enzyme located at the end of the poly(A) tail. The linearized plasmids were purified using the Nucleic Acid Purification Kit (Monarch). To generate the IVT mRNAs, we employed the HiScribe® T7 ARCA mRNA kit (NEB). The resulting mRNAs were purified using the Monarch RNA cleanup kit (NEB). Following nanodrop analysis of the mRNAs, isopropanol precipitation [28] was performed if any inorganic contaminants affecting the A230/260 ratio were detected. This involved mixing the mRNA samples with an appropriate amount of 3 M sodium acetate solution (pH 5.2) to achieve a final concentration of 0.3 M, followed by adding an equal volume of isopropanol. The mixture was then incubated at −20°C overnight. After incubation, the mRNAs were recovered by centrifugation at 14,000 g for 10 minutes. The resulting pellet was washed with 500 µl of ice-cold 70% ethanol and centrifuged under the same conditions. Any remaining ethanol was evaporated, and nuclease-free water was used to dissolve the mRNA samples. These samples were then stored at −80°C until the transfection experiments were conducted.

#### Cell line maintenance

We utilized the HEK293T and MCF7 mammalian cell lines for subsequent experiments. To prepare a complete growth medium for both cell lines, 440 ml of high glucose Dulbecco’s Modified Eagle’s Medium (DMEM) was mixed with 50 ml of heat-inactivated Fetal Bovine Serum (FBS), 5 ml of 100x L-Glutamine (200 mM), and 5 ml of 100x Penicillin/Streptomycin. This mixture was then filtered using a Corning Disposable Filter Unit with a pore size of 0.45 µm. The resulting medium was stored at +4°C. Cell media were changed every other day, and subculturing of both cell lines was performed when they reached 90% confluency.

#### mRNA transfection and sample collection

For the production of human TNF-*α* under different UTRs, mRNA transfection was performed on HEK293T cells. These cells were seeded into 6-well plates at a density of 300,000 cells per well, with each well containing 1 ml of growth medium. Following a 24-hour incubation period in a 37°C humidified incubator with 5% CO2, the media were aspirated, and 1 ml of fresh medium was added to each well. Subsequently, 5 µg of IVT mRNAs containing the 5’ UTRs (including human *β*-globin, UTR2, UTR5, UTR7, UTR9, and UTR10) were transfected into the cells using Lipofectamine 3000 (Invitrogen). One well served as a negative control without mRNA to assess the impact of the transfection reagent. The transfected cells were then returned to the incubator for an additional 24 hours. After the incubation, 1.2 ml of supernatant from each well was collected into ice-chilled tubes and kept on ice. To eliminate cellular debris, all samples were centrifuged at 5800 rpm for 5 minutes. The supernatants were concentrated to 400 µl using 0.5 ml 3 kDa centrifugal filters (Amicon) and stored at +4°C for subsequent cytotoxicity experiments, which were planned to be carried out on the same day to prevent degradation.

#### TNF-*α* cytotoxicity assay

The cytotoxic effect of TNF-*α* on cancer cells [70] was evaluated using the previously described methodology [25]. MCF7 cells were seeded into a 96-well plate at a density of 6,000 cells per well, with each well initially containing 100 µl of medium. After a 24-hour incubation period in a 37°C humidified incubator with 5% CO2, the old media were removed, and 65 µl of fresh medium was added to each well. This fresh medium was then combined with 35 µl of the concentrated TNF-*α* samples obtained from mRNA transfection. Top of Form All samples, including the negative control and those with different 5’ UTRs (such as human *β*-globin, UTR2, UTR5, UTR7, UTR9, and UTR10), were studied in triplicate. After 48 hours of incubation, the supernatants were removed from the wells, and 90 µl of fresh medium was added to each well. Subsequently, 10 µl of a 5 mg/ml MTT solution was added to the wells, and the plate was incubated for 4 hours at 37°C. Following this incubation, the supernatants were discarded, and 100 µl of DMSO was added to each well to dissolve the formazan crystals. The plate, wrapped in aluminum foil, was then shaken for 15 minutes to ensure complete dissolution of the crystals. The absorbance at 570 nm was measured for each well, and the viability of each treatment was calculated by comparing it with the negative control sample.

## Competing Interests

The authors declare no competing interests.

## Author Contributions

SB and FO designed and implemented the computational model. SB, AH, UOSS, and AEC wrote the manuscript. UOSS and AEC supervised the study.

## Data Availability

All the data used here are available in public repositories. We downloaded the dataset from the UTRdb 2.0 database [41]. The Human gene sequences are exported from Ensemble Biomart [18].

## Code Availability

The source code, including the implementation of the model, the optimization procedure, and the additional scripts used to generate plots are released at http://github.com/ciceklab/UTRGAN.

## Notes

### Competing Interest Statement

The authors have declared no competing interest.

### Summary of Updates

We have done further analysis and included more motif-based comparisons of the sequences, as well as biologically meaningful analysis of the sequences.

