## Supplemental Material for "UTRGAN: Learning to Generate 5’ UTR Sequences for Optimized Translation Efficiency and Gene Expression"

#### 1 Supplementary Figures

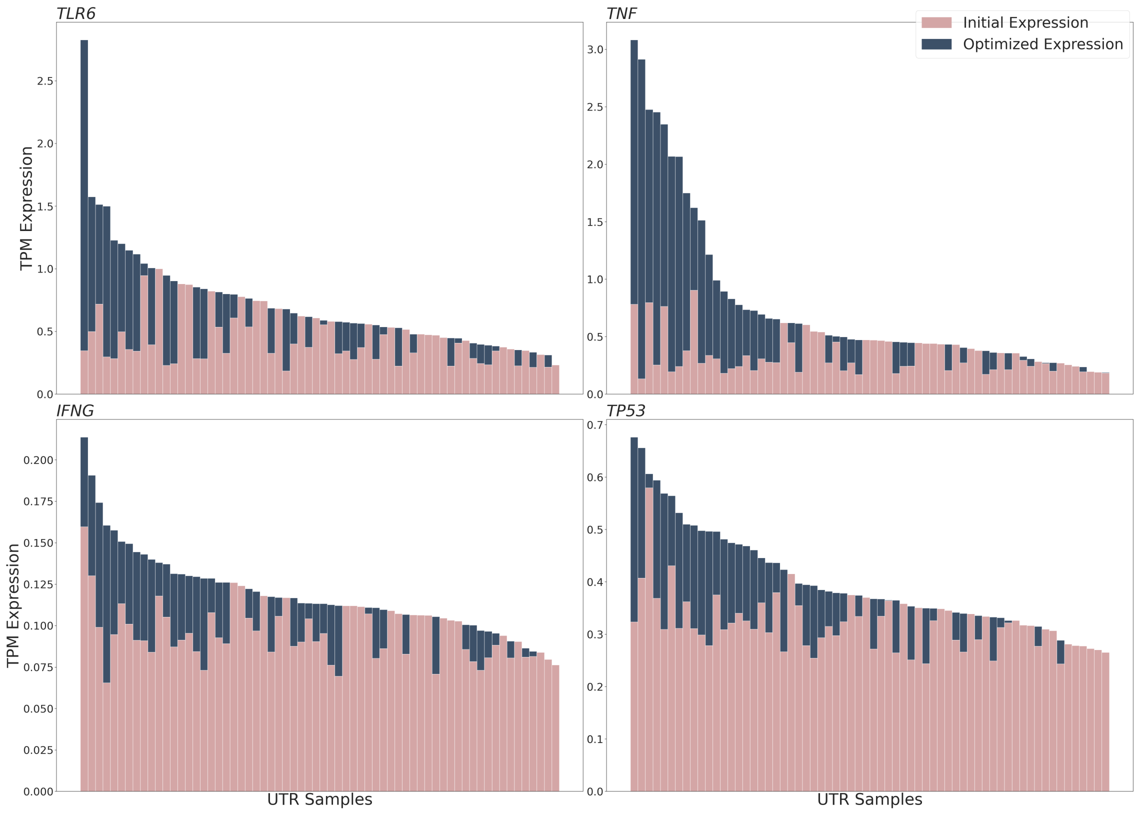


Supplementary Figure 1: **Optimization with maximum GC content limitation.** The optimized sequences with a maximum GC content limitation of 65%. We performed the optimization and selected the best sequence with the condition of their GC content being below the selected value. Although this method does not perform as well as the uncontrolled optimization in increasing the gene expression, it does result in considerable increases in all four genes (*TLR6*, *IFNG*, *TNF*, *TP53*) while maintaining the GC content. The optimized expression values are behind the initial values in all panels, and for the few sequences where the blue bar is not shown, the optimized value is smaller than the initial value.

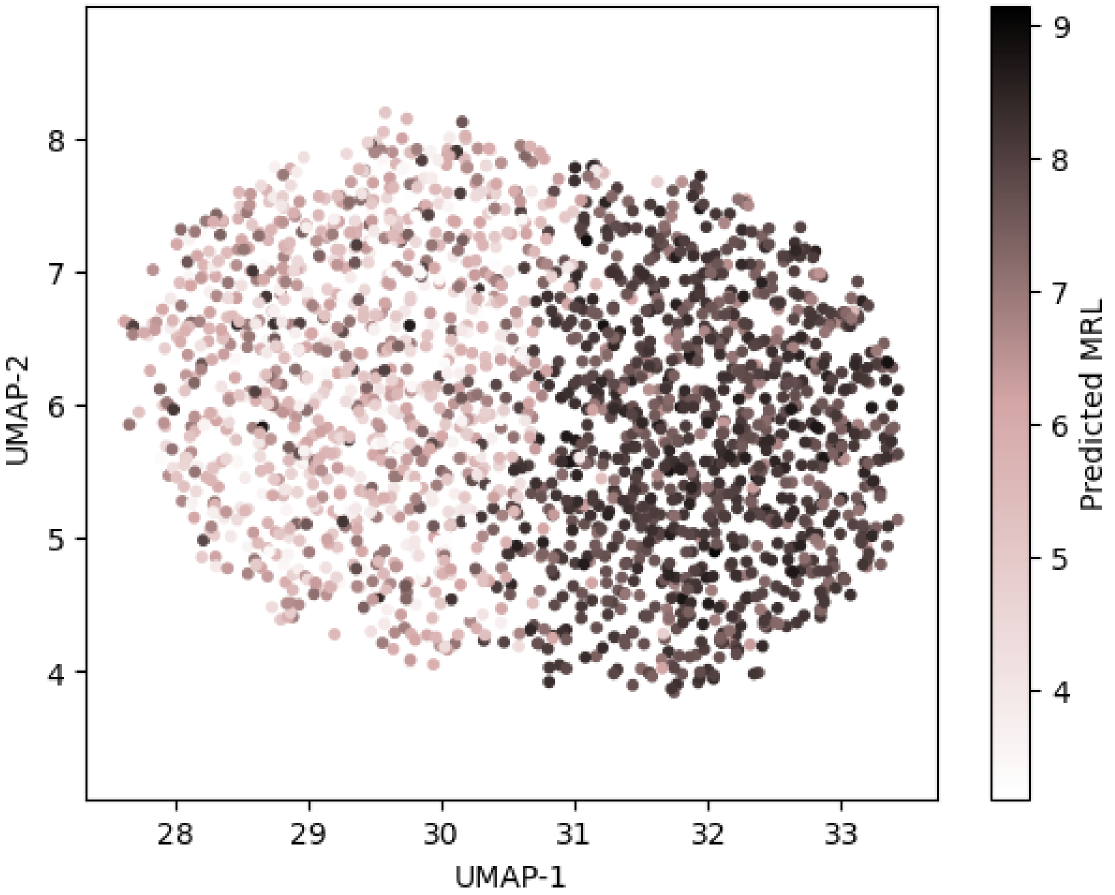


Supplementary Figure 2: **The latent space of UTRGAN for generated and optimized UTRs.** Here, we show the latent space of the model for 5' UTRs generated and optimized using UTRGAN. The UMAP is performed on the latent vector of 2048 sequences, including 1024 initially generated sequences and 1024 sequences resulting from optimization of the first set of sequences. The reduced vectors are colored by their predicted MRL values. The plot shows that the model has learned a meaningful latent space, and sequences before and after optimization are well-separated in the latent space.

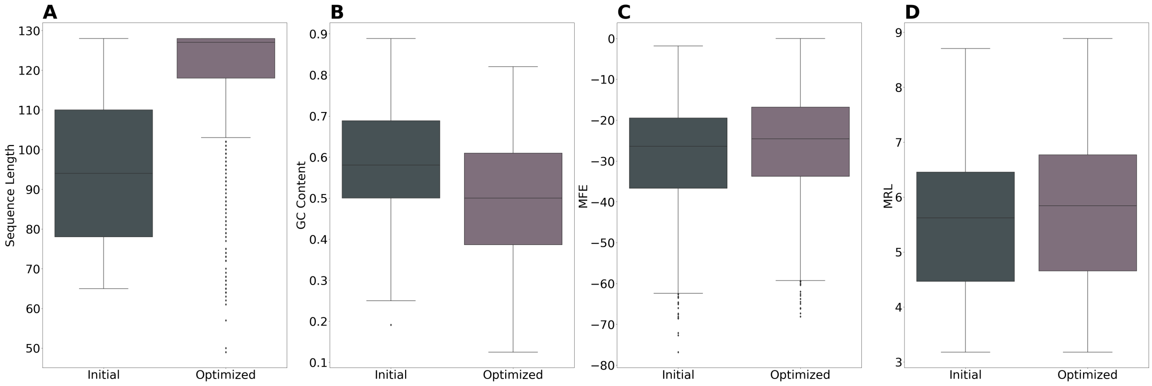


Supplementary Figure 3: **Effect of TE optimization with respect to length, GC content, MFE, and MRL of the sequences. The box plots characterize the values using the 25th, 50th, and 75th, also known as quartiles (Q1, median, Q3) and the interquartile range (IQR = Q3 - Q1), with whiskers extending to a maximum of 1.5 times the IQR. Outliers beyond the whiskers are plotted separately.**

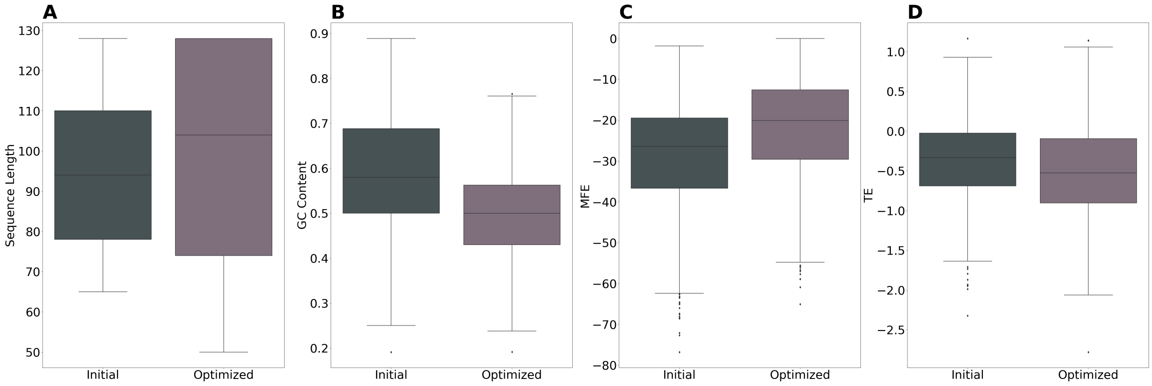


Supplementary Figure 4: **Effect of MRL optimization with respect to length, GC content, MFE, and TE of the sequences. The box plots characterize the values using the 25th, 50th, and 75th, also known as quartiles (Q1, median, Q3) and the interquartile range (IQR = Q3 - Q1), with whiskers extending to a maximum of 1.5 times the IQR. Outliers beyond the whiskers are plotted separately.**

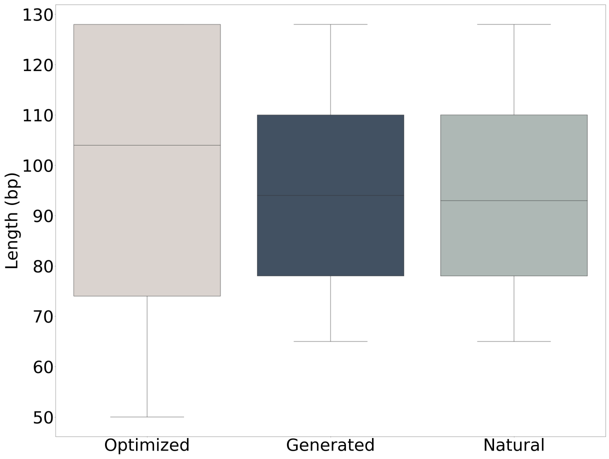


Supplementary Figure 5: **Distribution of length of the generated, MRL-optimized, and natural 5' UTRs**

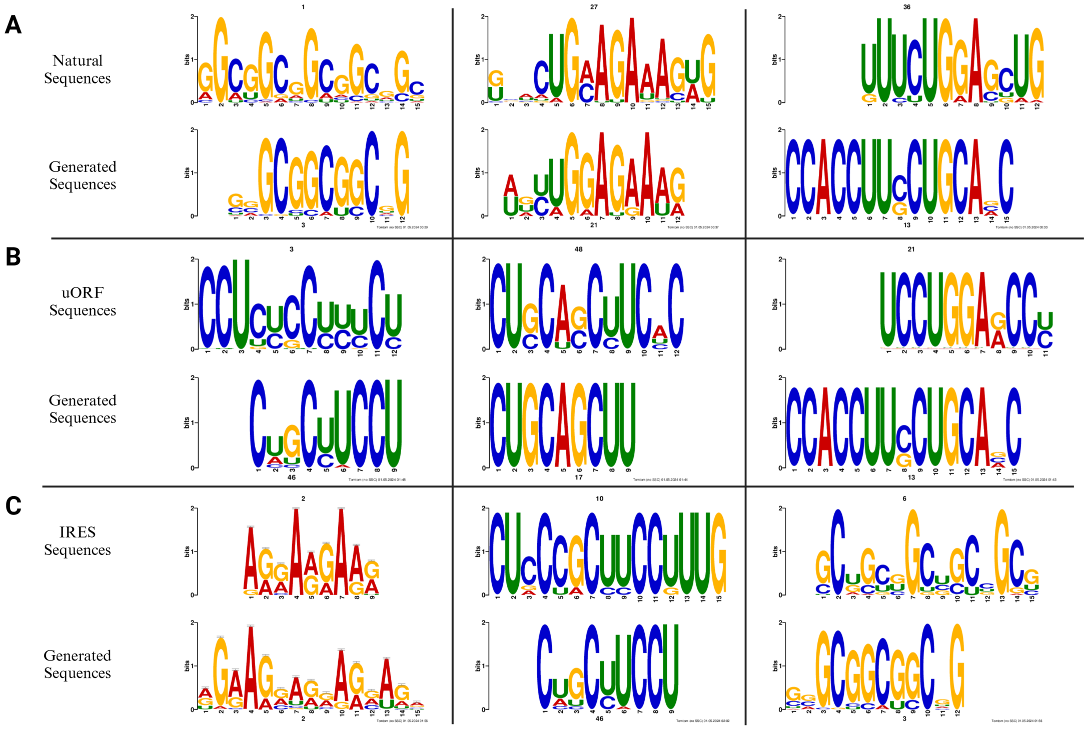


Supplementary Figure 6: **Top Matching Motives between UTRGAN-generated and various natural sequences.** The top-3 motif matches between the sequences optimized using UTRGAN and the natural 5' UTR, uORF, and IRES sequences are shown on panels **A**,**B**, and **C**, respectively. The leftmost motif is the top match for each category.

### 2 Supplementary Notes

#### 2.1 Experimental Setup

The model is implemented using the Tensorflow 2 library and in Python. We train and optimize the model on a Super-Micro Super-Server 4029GP-TRT with 2 Intel Xeon Gold 6140 Processors (2.3 GHz, 24.75 M cache), 256 GB RAM, 6 NVIDIA GeForce RTX 2080 Ti GPUs (11 GB, 352 bit), and 2 NVIDIA TITAN RTX GPUs (24 GB, 384 bit). We only use a single TITAN RTX GPU. The training takes around 14 hours for 3,000 epochs, and we select this checkpoint based on the validation loss as well as the $p$-values obtained from the statistical tests. The TE, MRL, and gene expression optimization take 3, 4, and 15 minutes, respectively. Optimizing UTR sequences for higher average expression in 8 genes takes 90 minutes.

#### 2.2 Hyperparameter Optimization

The main hyperparameters of our model include the size of the latent dimension, the number of generator layers, and the number of discriminator layers. To find the best combination of parameters, we run a grid search on a reasonable range of parameters. The model is trained up to 200,000 generator iterations. Many of the instances of the model fail to learn and overfit, resulting in the validation loss going below the training loss, and the training loss begins to increase in some cases. In the cases that the mentioned scenario doesn't happen, we select the parameters where the $p$-values for the statistical tests are closer to desirable values. The latent dimension in UTRGAN is 40, and we present the number of layers in Supplementary Tables 6 and 7.

#### 3 Supplementary Tables

**Supplementary Table 1.** The top generated 5' UTR sequences for high gene expression in the target genes, that are *TLR6*, *IFNG*, *TNF*, and *TP53*. The table shows the top 5' UTR for each gene. These sequences result in higher predicted expression than the other generated ones when attached to these genes as 5' UTRs.

| Gene | 5' UTR |
|---|---|
| TLR6 | CGGUGUCUGGAAGCCUUCUUUCGCGGUGCUGGACUCCGGGCCCGA<br>GGCCCCGAAUAGUCGCGGAAGUUCUGGGACGAAUCCGGGGCCCGG<br>UCGAACCGGCGGGGGCGCCGGCCGGCCGUCCAG |
| IFNG | AGACGUGGACCGCUUCGCGGCGGCUGUCGGGUGGAGUGGGAGGCG<br>AGCCGGAAGUGGCGGUCCUGGCUGCUCCGGGUCGUCCCUGCCGGC<br>GUCGGAGCGUCCCUCGGAACCAGGAGUGCCGGCGACG |
| TNF | GAGUUGCGGCCGGGCGGCGUCGCGGCGGUCGGGGCCGAGAGUCGU<br>GGCCGGAAGCCGCCGUUGGAUGGGGCCGGGUCGGUUGGAGACGCC<br>GGCGGCCCCGCCGCCCCCTTTGGUGGGCCCGA |
| TP53 | CGGCUUGCUGCCGCCCGCGACAGCGGAGACAAGCAGGGAGGUCCU<br>GCCGCCUCCCAGAAAUAUUGGGGGGGCUGGGGAAGCCGCAGGCCG<br>ACUGAGUCGGGAGGCGGACGGCGGAAGGACGCGGCGGA |

**Supplementary Table 2.** The top generated 5' UTR sequences for joint optimization of translation efficiency and high gene expression in the target genes, that are *TLR6*, *IFNG*, *TNF*, and *TP53*. The table shows the top 5' UTR for each gene. These sequences result in higher predicted expression as well as higher predicted translation efficiency than the other generated ones when attached to these genes as 5' UTRs.

| Gene | 5' UTR |
|---|---|
| TLR6 | UGGCUGUGGCGGCCGGAAAUCUGAGCCUCCGGGCCAGUGGGCCGGG<br>AGCGGUCGGGUCUGGAGCGGCGGAGGGAGAACGGAAGCUGCGGAGG<br>CCCCGCACGCGGGGGGCUGAGGUCGGCCUCGUCAGU |
| IFNG | UGGCUGUGGCGGCAGGAAUUCUGUGCCUCCGGGCCUGUGGGUCGGG<br>AGCGGCCGGGUCUGGAGCGGCGGUGGGAGAAAGGAAGCGGCGGUGG<br>CCCCGCAAGCGGGGGGCUGAGGGCGGCCGGGGCGG |
| TNF | AGUCCGCUGGCGUGGCCCAGCCCCGGGGCCGGUUCUCGGGAACGAG<br>AAGCUGGUGAUCUCCGUUGGGGCGCGGUCGGAUGCGGUUGAGGGAG<br>GCGAGGACGCGAAAGCGGGCCCGCAGCGGAGCCGGC |
| TP53 | GGGAGGUGGUUCCGAAAUCGCCGUGAAGCCAAGCCCUCUGCUCCGU<br>UUCCUCUUUUCCACGGACUGGGAGGUUUUCGGGACCACGCCAGGAG<br>AACCACGAGGGCUAGGCAGAGGACUGAAGCGUCGGA |

**Supplementary Table 3.** The top generated 5' UTR sequences for GC content controlled high gene expression in the target genes, that are *TLR6*, *IFNG*, *TNF*, and *TP53*. The table shows the top 5' UTR for each gene. These sequences result in higher predicted expression compared to other generated 5' UTRs with GC content lower than 65%.

| Gene | 5' UTR |
|---|---|
| TLR6 | GUGAAAGGCGUGCCGCCGCCCUAGAGUCCUGGGAAGACAGUCAAG CGGAGAGAGGGAAGUUUCCCUGUACUUCCUCCCUCCCGGCCGCGAGU CCUGGCAUCCCGCUGAGGCUGG |
| IFNG | UGGCUGUGGCGGCAGGAAUUCUGUGCCUCCGGGCCUGUGGGUCGGG AGCGGCCGGGUCUGGAGCGGCGGUGGGAGAAAGGAAGCGGCGGUGG CCCCGCAAGCGGGGGGCUGAGGGCGGCCGGGGCGG |
| TNF | CUUCGCCCUUCUUCAUAAUCAGUCGACAAGCGGUUUCUGGAAGGCG GCGUCUCGCUGUUGCUGUGGCCUCUGGAAGCAGGAGAAGAGGCGGC UUGGAAACGCUGGAAGCGGCCGCGGCGUCCUCAG |
| TP53 | AAGACGAGGCCAGGCUGCGCAAAUCCGGCUGGGGCCGAGUUCUUGG UCCGUUUUCCCAAUAAGAGUGGUCCCGGGAAUGGACGAGACGCCGC CGGAAGCGUCUCUGGAGGUUGAUGGCAAG |

**Supplementary Table 4.** The top 10 generated 5' UTR sequences for optimized MRL.

| UTR | Predicted MRL | UTR |
|---|---|---|
| GAGGAGCAGAAAUUGGCCGCUGUUGCGUUAAUCCACAACGGUGCUGUUG UGAGUUGACAGCAGGUCCAGGGAUAAGGUGUGGUGAAGUUGGUGUCUGUAG CAUAUCCAUGGAGAUGAUUGCAAUAGUA | 9.06 | UTR1 |
| UUUAACACAUAUCAGAAAAAGAAUCAAGAACUGUUGGAGCCGGGUUAAAA AAAACUUUCCAAGUAAUGUUAAAG | 9.02 | UTR2 |
| CUUUCCUGAGAAUUCCCUGGAAGCACUUGUUCAACCACAGCACUAGGUUUC AAAGCAAGGUUCACACCCAGCCUCCACAAGGCCUGGGGAUGAUGAAGAGG GGAAUA | 8.91 | UTR3 |
| GUUUUAUCCAGCAUCCAUUCGAGAACUUGAUCAAGUCCCACAUGCUACCCAG GACUAGGAGAGAAGGAAGCAGAAAGACAGUAAAUUGCCGCGGAUAUUUUCAG | 8.81 | UTR4 |
| GUUACCAUACUGUAAUCAGAAUUCUGCUUCUAAGGAUAAAUUUACUUCUACU UAUUAUAAAAAAAG | 8.80 | UTR5 |
| UGGACUCAGUCCCAACCAAAUAGUGGUGUUUCUGCAGCUCCAAACUUGAGAGA AGAACUUCAAGCUCUCCUUUAAACCGGUGUUUGC | 8.74 | UTR6 |
| GUGUUUCCCAUUUAGUCACUAAGAGGUGACGCCAUACCACAUACAGGUGGAGG AGCAUUAAAGC | 8.577 | UTR7 |
| GGGUGGUCGGAGCGGCUGCAGGAGGCAGUAGGAAGAAGAGAAAAGGAAGGAGG AGGAUCCGAACACGUCACAAAACAGUCGG | 8.575 | UTR8 |
| AGAAACCAGAAGACCUUCUGGAGGAAGACUUC CAUAGCUUUGUCUCAGUGAAGGCAGC | 8.56 | UTR9 |
| AGAAAUUUCAGUAAUAAUCGUCCUGGAGGAGUGGGUAACAUAUUUGAAGAUUUC CAUUUCAGCUUU | 8.49 | UTR10 |

**Supplementary Table 5.** The top 3 generated 5' UTR sequences for optimized translation efficiency. The predicted *log*-TE for these sequences is 1.5, 1.3, and 1.2, respectively.

| |
|---|
| GAAUCUCAUCUUCGCCCUUAACCAUCUCAGAGACCUUCCAACAGAA<br>CUUGGUGUAAAAGCCAGGCUUCCCCGCCUCUGCAGUGCUGCGGCUU<br>GGCUGCCUCUCCGCGCGAGG |
| GACCAAGCACACCUCACCCCGCGCCCCCAGUCCGGGCGCCGGGCUC<br>CUCGGUGGUCUCAGCUGCU |
| ACCAUCCCUCCAGCAGCUGGGACGUGCCUGCGCCGCAGCCGUGGCC<br>GCCUCGCUGCGGCUGUCUCCCCCCAG |

**Supplementary Table 6.** The layers of the Generator in the WGAN model we use to generate realistic 5' UTR sequences. In this configuration, the batch size for training the model is 64, as seen in the table.

| Input/Layer | Kernel size | Output shape |
|---|---|---|
| $z_0$ | - | 64 x 40 |
| Dense | - | 64 x 5120 |
| Reshape | - | 64 x 128 x 40 |
| Convolution (1D) | 1 | 64 x 128 x 40 |
| Residual block (1D) | [5]x2 | 64 x 128 x 40 |
| Residual block (1D) | [5]x2 | 64 x 128 x 40 |
| Residual block (1D) | [5]x2 | 64 x 128 x 40 |
| Residual block (1D) | [5]x2 | 64 x 128 x 40 |
| Residual block (1D) | [5]x2 | 64 x 128 x 40 |
| Convolution (1D) | 1 | 64 x 128 x 5 |
| Softmax | - | 64 x 128 x 5 |

**Supplementary Table 7.** The layers of the Discriminator (Critic) in the WGAN model we use to generate realistic 5' UTR sequences.

| Input/Layer | Kernel size | Output shape |
|---|---|---|
| Convolution (1D) | 1 | 64 x 128 x 40 |
| Residual block (1D) | [5]x2 | 64 x 128 x 40 |
| Residual block (1D) | [5]x2 | 64 x 128 x 40 |
| Residual block (1D) | [5]x2 | 64 x 128 x 40 |
| Residual block (1D) | [5]x2 | 64 x 128 x 40 |
| Residual block (1D) | [5]x2 | 64 x 128 x 40 |
| Flatten | - | 64 x 5120 |
| Dense | - | 64 x 1 |
